# Pre-clinical Evaluation of Biomarkers for Early Detection of Nephrotoxicity Following Alpha-particle Radioligand Therapy

**DOI:** 10.1101/2023.09.27.559789

**Authors:** Mengshi Li, Claudia Robles-Planells, Dijie Liu, Stephen A. Graves, Gabriela Vasquez-Martinez, Gabriel Mayoral-Andrade, Dongyoul Lee, Prerna Rastogi, Brenna M. Marks, Edwin A. Sagastume, Robert M. Weiss, Sarah C. Linn-Peirano, Frances L. Johnson, Michael K. Schultz, Diana Zepeda-Orozco

## Abstract

**Purpose:** Cancer treatment with alpha-emitter-based radioligand therapies (α-RLTs) demonstrates promising tumor responses. Radiolabeled peptides are filtered through glomeruli, followed by potential reabsorption of a fraction by proximal tubules, which may cause acute kidney injury (AKI) and chronic kidney disease (CKD). Because tubular cells are considered the primary site of radiopeptides’ renal reabsorption and potential injury, the current use of kidney biomarkers of glomerular functional loss limits the evaluation of possible nephrotoxicity and its early detection. This study aimed to investigate whether urinary secretion of tubular injury biomarkers could be used as additional non-invasive sensitive diagnostic tool to identify unrecognizable tubular damage and risk of long-term α-RLTs nephrotoxicity.

**Methods:** A bifunctional cyclic peptide, melanocortin ligand-1(MC1L), labeled with [^203^Pb]Pb-MC1L, was used for [^212^Pb]Pb-MC1L biodistribution and absorbed dose measurements in CD-1 Elite mice. Mice were treated with [^212^Pb]Pb-MC1L in a dose escalation study up to levels of radioactivity intended to induce kidney injury. The approach enabled prospective kidney functional and injury biomarker evaluation and late kidney histological analysis to validate these biomarkers.

**Results:** Biodistribution analysis identified [^212^Pb]Pb-MC1L reabsorption in kidneys with a dose deposition of 2.8, 8.9, and 20 Gy for 0.9, 3.0, and 6.7 MBq injected [^212^Pb]Pb-MC1L doses, respectively. As expected, mice receiving 6.7 MBq had significant weight loss and CKD evidence based on serum creatinine, cystatin C, and kidney histological alterations 28 weeks after treatment. A dose-dependent urinary Neutrophil gelatinase-associated lipocalin (NGAL, tubular injury biomarker) urinary excretion the day after [^212^Pb]Pb-MC1L treatment highly correlated with the severity of late tubulointerstitial injury and histological findings.

**Conclusion:** urine NGAL secretion could be a potential early diagnostic tool to identify unrecognized tubular damage and predict long-term α-RLT-related nephrotoxicity.

## Introduction

Radioligand therapies (RLTs) have emerged as a form of therapy in oncology that has the potential to be transformative for cancer patients. These radiopharmaceuticals consist of a radionuclide coupled to a small peptide ligand that binds to cell membrane targets with high affinity and specificity, making these agents attractive as cancer therapeutics (**Figure 1a**). Unlike external beam radiation therapy (EBRT), these targeted forms of systemic radiotherapy enable radionuclides to be delivered directly to cancer cells. Examples of FDA-approved radiopharmaceuticals that are significantly improving outcomes for cancer patients include beta (β)-particle emitting RLTs LUTATHERA^®^ (*i.e.*, [^177^Lu]Lu-DOTATATE) for advanced neuroendocrine tumors (NETs) and PLUVICTO^®^ (*i.e.*, [^177^Lu]Lu-PSMA-617) for metastatic castrate-resistant prostate cancer (mCRPC) targeting somatostatin receptor type 2 (SSTR2) and prostate-specific membrane antigen (PSMA), respectively. RLTs are also effective in other SSTR-expressing tumors, such as paraganglioma and pheochromocytoma, thyroid carcinoma, and meningioma[1]. Use of other ligands for overexpressed receptor targets has been demonstrated in breast, colon, myeloma, and ovarian cancer[1], melanoma[2], and hematological malignancies[3]. There is a significant difference in the observed efficacy of RLTs that is dependent on the radionuclide utilized. Compared to FDA-approved β-emitter bioldrugs, recent studies using alpha (α) emitters-RLTs (*e.g.*, ^212^Pb, ^225^Ac, ^211^At) in patients with NET and mCRPC, demonstrate greater tumor cell cytotoxicity, including patients with refractory disease following β-RLT[4]. Fundamentally, the advantages of α-emitters lie in the higher linear-energy transfer (LET) (∼100 keV/µm), which induces double-strand DNA breaks (DSB) and a concomitant increase in ionizations (primary and secondary) along their short path length in the cellular tissue microenvironment[5]. The deposition of energy over extremely short path length results in a significantly increased incidence of lethal DSB; high relative biological effectiveness (RBE); improved tumor-cell-cytotoxicity; and activation of anti-tumor immunity [6]. Among commercially available α-emitters, [^212^Pb] (half-life T_1/2_ 10.6 h) is produced using a [^224^Ra]/[^212^Pb] generator system that enables nimble on-demand production of [^212^Pb]Pb-based radiopharmaceuticals[7]. In the case of [^212^Pb], the isotope [^203^Pb] (T_1/2_ 51.9 h) can be used as a single-photon emission computerized tomography (SPECT) imaging diagnostic companion to provide detailed dosimetry that guides [^212^Pb] therapy dosing [8]. The [^203^Pb]/[^212^Pb] radionuclide pair has the advantage that elementally identical imaging surrogate [^203^Pb] can guide [^212^Pb] therapy (**Figure 1b**). Here, we establish biomarkers that can be used in the preclinical setting to connect surrogate imaging information to the potential for acute kidney injury (AKI) to more effectively use the imaging data to establish appropriate therapeutic dosing strategies. It is envisioned that these data can lead not only to more effective dose planning for clinical translation of new RLTs but for a broader spectrum of pharmaceutical drug candidates for clinical introduction for cancer therapy.

**Fig 1.**
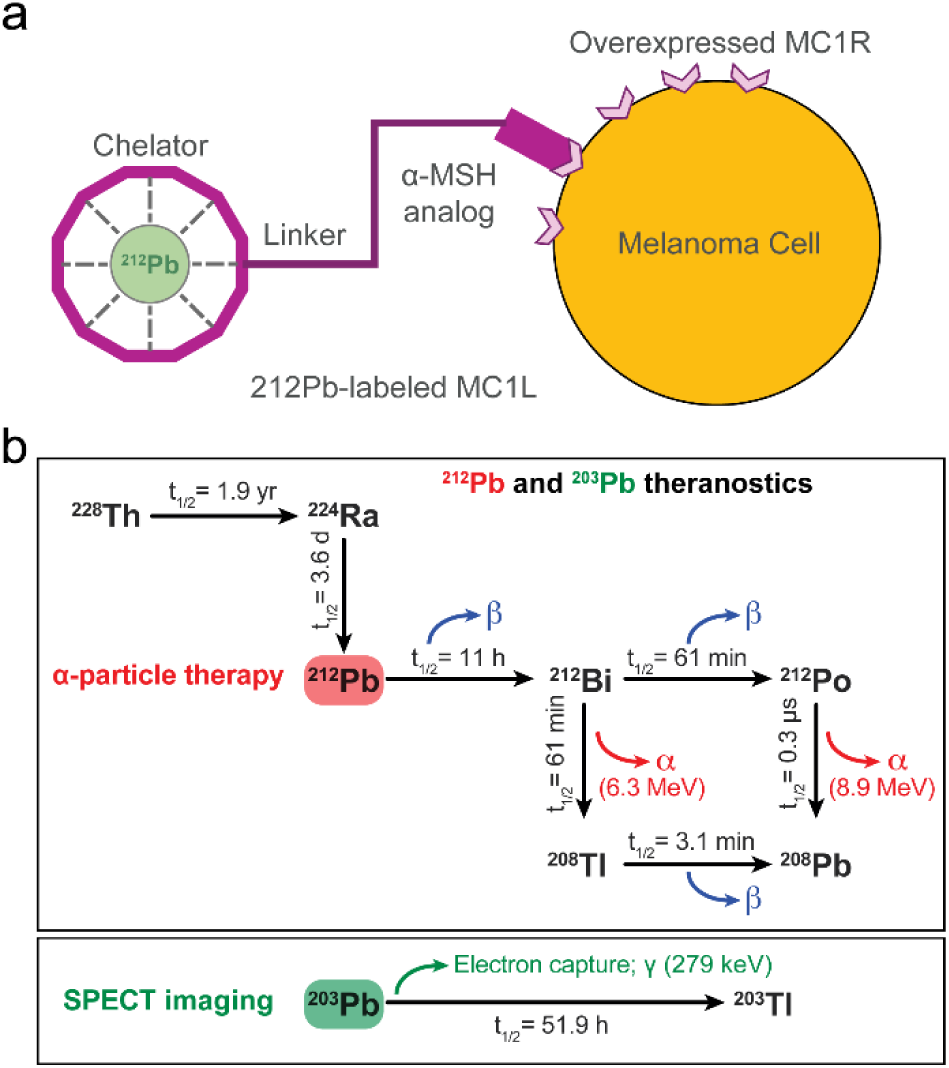
(a) Schematic representation of [^212^Pb]Pb-labeled melanocortin 1 ligand (MC1L), melanoma cell expressing melanocortin 1 receptor (MC1R), and alpha melanocyte stimulating hormone (α-MSH) analog. (b) Schematic representation of [^212^Pb] and [^203^Pb] nuclides decay.

While emerging research demonstrates the potentially transformative impact of α-particle RLT for cancer treatment, kidneys are often treatment-limiting due to their role in bioelimination of unbound therapeutic agents. While agents can be pharmacologically designed to minimize the reabsorption, biomarkers that provide information on the potential for nephrotoxicity are needed because current evaluation is limited to late post-therapy histological findings and estimation of glomerular filtration rate (eGFR) using the endogenous maker of kidney function serum creatinine (sCr). Fortunately, due to their relatively small size (< 5.5 kDa) and hydrophilic nature, radiolabeled peptides are rapidly filtered through glomeruli. However, a fraction can be reabsorbed in tubular cells, which may lead to injury[9]. Studies in tubule-specific megalin knockout mice demonstrate that proximal tubules are primarily, but not exclusively, responsible for radiopeptide uptake within the kidneys[10]. Because of the short travel distance of emitted particles (low-energy β<1.7 mm^1^; α<80 µm), the radiation exposure from RLT, especially α-RLT, may be highly localized in the proximal tubular area rather than glomeruli. This suggestion is further supported by prior pre-clinical studies evaluating nephrotoxicity of alpha and beta-emitter RLT that demonstrated the presence of tubulointerstitial disease as early as 10 weeks, without significant glomerular changes, which progressively worsens and leads to severe chronic kidney disease (CKD) seven months after therapy[11,12]. This observation is consistent with the known pathophysiology of AKI-induced CKD, wherein primary tubular cells that fail to repair after severe or recurrent injuries adopt a pro-fibrotic and pro-inflammatory phenotype that drives progressive kidney damage resulting in structural abnormalities including fibrosis, vascular rarefaction, tubular loss, glomerulosclerosis, and a chronic inflammatory infiltrate, with subsequent renal function decline[13]. Thus, primary tubular damage alone could be the etiology of RLT-induced long-term kidney toxicity.

Current nephrotoxicity screening for assessing pharmaceutical-treated patients and mice for establishing potential kidney injury depends mainly on prospective sCr measurement and its routine use to calculate eGFR. Kidney toxicity in patients receiving RLT is expected to be associated with patchy loss of tubule cells, focal areas of proximal tubular dilation, and distal tubular casts, together with measurements of cellular regeneration[14]. Serum creatinine is a delayed and insensitive biomarker of kidney function decline that does not reflect the degree of tubulointerstitial damage. Additionally, long-term toxicity evaluation is limited by the patient population’s poor survival. In comparison, urinary biomarkers such as Neutrophil Gelatinase-Associated Lipocalin (NGAL), Kidney Injury Molecule 1 (KIM-1), and Epidermal Growth Factor (EGF) have been used to detect AKI early in patients with normal sCr (subclinical AKI) and to evaluate overall tubular health[15]. The change in their urinary secretion pattern has been used to identify patients at risk of progressive CKD[16]. Predictive biomarkers could accurately identify patients with subclinical kidney injury and those at risk of AKI and CKD following RLT treatment.

Regulatory bodies currently employ 23 Gy in kidneys as the absorbed dose limit, derived from EBRT, despite significant differences in the form and interaction of radioactivity emissions associated with RLT vs. EBRT. For example, the typical dose rate for EBRT is ∼1 Gy/min, whereas the absorbed dose from RLT is delivered over multiple days, even weeks, depending on the half-life of the radionuclide, resulting in a slower dose rate (mGy/min). Although conventional EBRT dose limits may differ from those encountered with RLT, dosimetry-based therapy remains a potentially powerful tool for personalizing RLT. Specifically, in RLT a biological effective dose of 40 Gy to the kidney has been suggested to be safe in patients without risk factors for kidney disease and 28 Gy in those at risk of kidney injury[17]. A more precise understanding of the nature of kidney-delivered doses may help maximize cancer therapy and reduce long-term kidney toxicity according to individual patient tolerability. However, this approach is limited by the need for more information on tubular damage caused by a specific kidney-absorbed dose not provided by sCr measurement. Thus, the purpose of this study was to investigate the urinary secretion pattern of tubular injury biomarkers following injection of a [^212^Pb]Pb-based RLT in a murine model. In this dose-escalation study, a bifunctional cyclic peptide, melanocortin 1 ligand (MC1L), was labeled with [^203^Pb]Pb-MC1L to determine the distribution and absorbed dose of [^212^Pb]Pb-MC1L at levels intended to induce AKI, followed by a comprehensive nephrotoxicity analysis including novel urine secretion pattern of kidney injury biomarkers in CD-1 Elite mice. To explore potential absorbed dose tolerated in humans, we performed human dosimetry scaling up from mice. It is envisioned that combining biomarkers and dosimetry can lead to a more detailed understanding of the use of [^203^Pb]-based imaging as a quantitative guide for [^212^Pb] therapy planning and a personalized approach to cancer therapy.

## Methods

### Reagents and Materials

Detailed information on reagents and material used in this study can be found in the **Supplemental Methods** section.

### Radiochemistry, *In Vivo* Biodistribution of [^203^Pb]Pb-MC1L and Dosimetry of [^212^Pb]Pb-MC1L

Radiochemistry radiolabeling and quality control of [^203^Pb]/[^212^Pb]Pb-MC1L were performed as previously described[6] and are detailed in **Supplemental Methods** section. Biodistribution of [^203^Pb]Pb-MC1L in normal organs/tissues was determined in male CD-1 Elite mice. All animal studies followed the Guide for the Care and Use of Laboratory Animals and were approved by the University of Iowa Animal Care and Use Committee. Mice were injected with 74 MBq [^203^Pb]Pb-MC1L (2-100 pmole injected peptide) in 100 µL saline *via* tail vein and were subsequently euthanized at 0.5, 1.5, 3, and 24 hours post-injection for organs/tissues resection (n=4-6 at each time point). Radioactivity in each organ was measured on Cobra II automated gamma counter and decay corrected. Data were normalized and expressed as percent injected dose per gram of tissue (%ID/g). The absorbed dose from [^212^Pb]Pb-MC1L injection in mouse and human were calculated using the [^203^Pb]Pb-MC1L surrogate biodistribution data using typical Medical Internal Radiation Dose (MIRD) absorbed fraction methods within the OLINDA/EXM v2.2 software. Detailed dosimetry calculations are described in **Supplemental Methods** section.

### Treatment to evaluate alpha-emitter toxicity

Male CD1-Elite mice were treated with a dose escalation regiment to evaluate toxicity at levels intended to cause kidney injury. Mice were administered either 0.0 (vehicle), 0.9, 3.0, and 6.7 MBq [^212^Pb]Pb-MC1L (injected peptide mass 0.0 – 0.6 nmole, n=6 per group) as single injection *via* tail vein on day 0. No lysine co-injection was conducted to ensure kidney damage induction by [^212^Pb]Pb-MC1L. Following treatments, body weight of individual animals was measured twice weekly during the acute stage (*i.e.*, the first 3 weeks post-injection) and once per week in later stages.

### Hematotoxicity, Blood Chemistry, and Cardiotoxicity

Hematotoxicity, liver toxicity, and cardiac structure and function were assessed as described in **Supplemental Methods** section.

### Urine Nephrotoxicity Biomarkers

Biomarkers of kidney injury and tubular health were measured in urine samples collected using metabolic cages at 1 week (day 1 and 3), 8 weeks, and 28 weeks (7 months) post-[^212^Pb]Pb-MC1L injection. Murine metabolic cages from Hatteras Instruments (Grantsboro,NC) were kindly provided by Dr. Jonathan Nizar at the Department of Internal Medicine at The University of Iowa. After collection, urine samples were kept on ice and centrifuged at 2000 rpm for 5 minutes to remove particulates, and the supernatant was stored in a −80°C freezer until the analysis day. Urine NGAL, KIM-1, EGF, and protein and creatinine levels were measured as detailed in **Supplemental Methods** section.

### Kidney Histological Analysis

Twenty-eight weeks post [^212^Pb]Pb-MC1L injection, mice were euthanized. Kidneys were fixed in 4% paraformaldehyde and embedded in paraffin. Tissue sections (5 microns thick) were stained with Period Acid-Shift (PAS) for tubular injury (TI), glomeruli changes (GC), and interstitial inflammation (IF) semiquantitative evaluation. To measure renal interstitial fibrosis, paraffin-embedded kidney sections were stained with Trichrome. A blinded pathologist performed scoring as described in **Supplemental Table 1**.

### Statistical and Correlation Analysis

Statistical analysis was conducted on GraphPad Prism 8 (GraphPad Software, San Diego, CA, USA). Two-way ANOVA was applied in CBC, serum chemistry, and serum biomarkers *p<0.05. Kruskal-Wallis test for histological score analysis, with * p<0.05. Urine biomarkers and body weight statistical analysis was made by two-way ANOVA mixed-effect analysis with * p<0.05. Correlation analysis was made by computed nonparametric Spearman correlation for urine biomarkers and multiparameter correlation.

## Results

### Biodistribution of [^203^Pb]Pb-MC1L and Dosimetry of [^212^Pb]Pb-MC1L

To determine [^212^Pb]Pb-MC1L absorbed dose in critical organs (*e.g.*, kidneys), we employed [^203^Pb]Pb-MC1L as a dosimetry surrogate in the *in vivo* biodistribution study in male CD-1 Elite mice. Following i.v. injection of 74 kBq [^203^Pb]Pb-MC1L, there was a rapid [^203^Pb]Pb-MC1L clearance through the kidneys and bladder (8.2%; **Figure 2a**). Within 1.5 hours post-injection, the remaining [^203^Pb]Pb-MC1L in blood was 0.06 %ID/g. The highest kidney accumulation (8.2%ID/g) was at 0.5 hours post-injection, with minimal accumulation in other organs. As intended, [^203^Pb]Pb-MC1L retention in kidneys was relatively low (1.9%ID/g at 24 hours) but higher than in other organs, resulting in higher radiation exposure, enabling our evaluation of biomarkers. We used this information to perform a dosimetry analysis within the MIRD schema to determine the absorbed dose in the kidneys resulting from administering [^212^Pb]Pb-MC1L in male CD-1 mice. Based on the OLINDA/EXM 2.2 35-gram mouse phantom, the average absorbed dose per injected activity in kidneys was 2.99 Gy/MBq, among which 92.7% resulted from α-radiation (**Figure 2b**), with an estimated tubular and glomerular dose per administered activity of 6.31 Gy/MBq and 16.3 Gy/MBq respectively (implementation of the Hobbs et al. model is detailed in Supplemental Material) [18]. Thus, the kidney-absorbed dose in male CD-1 mice delivered from 0.9, 3.0, and 6.7 MBq [^212^Pb]Pb-MC1L were 2.8, 8.9, and 20 Gy on average. Detailed murine dosimetry results are summarized in **Supplemental Table 5 and 6**. Finally, to determine a human absorbed dose from this therapy, we performed dosimetry in human subjects scaling up from mice as described in Supplemental Methods. Co-localizing progeny with parent [^212^Pb], the predicted human absorbed radiation dose in kidneys was 5.34 mGy/MBq, among which 4.76 mGy/MBq is contributed by α-radiation. Due to the rapid clearance and minimal accumulation in bone, the predicted human absorbed dose in red marrow was 0.26 mGy/MBq. Murine and human dosimetry results are summarized in **Figure 3** **and Supplemental Tables 6 and 7**, respectively.

**Fig 2.**
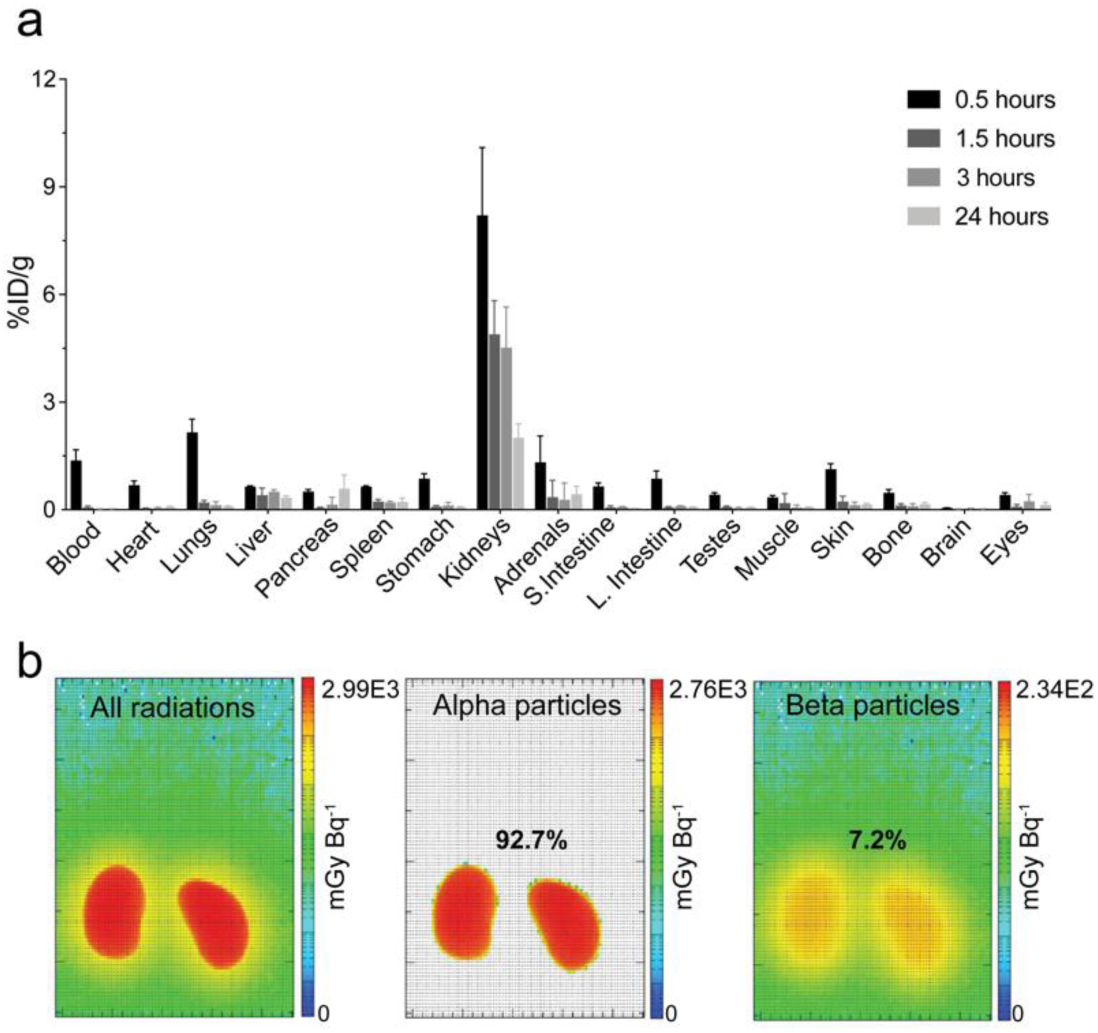
(a) Biodistribution of [^203^Pb]Pb-MC1L in male CD1 Elite mice. (b) Kidney dosimetry of [^212^Pb]Pb-MC1L in male CD1 Elite mice.

**Fig 3.**
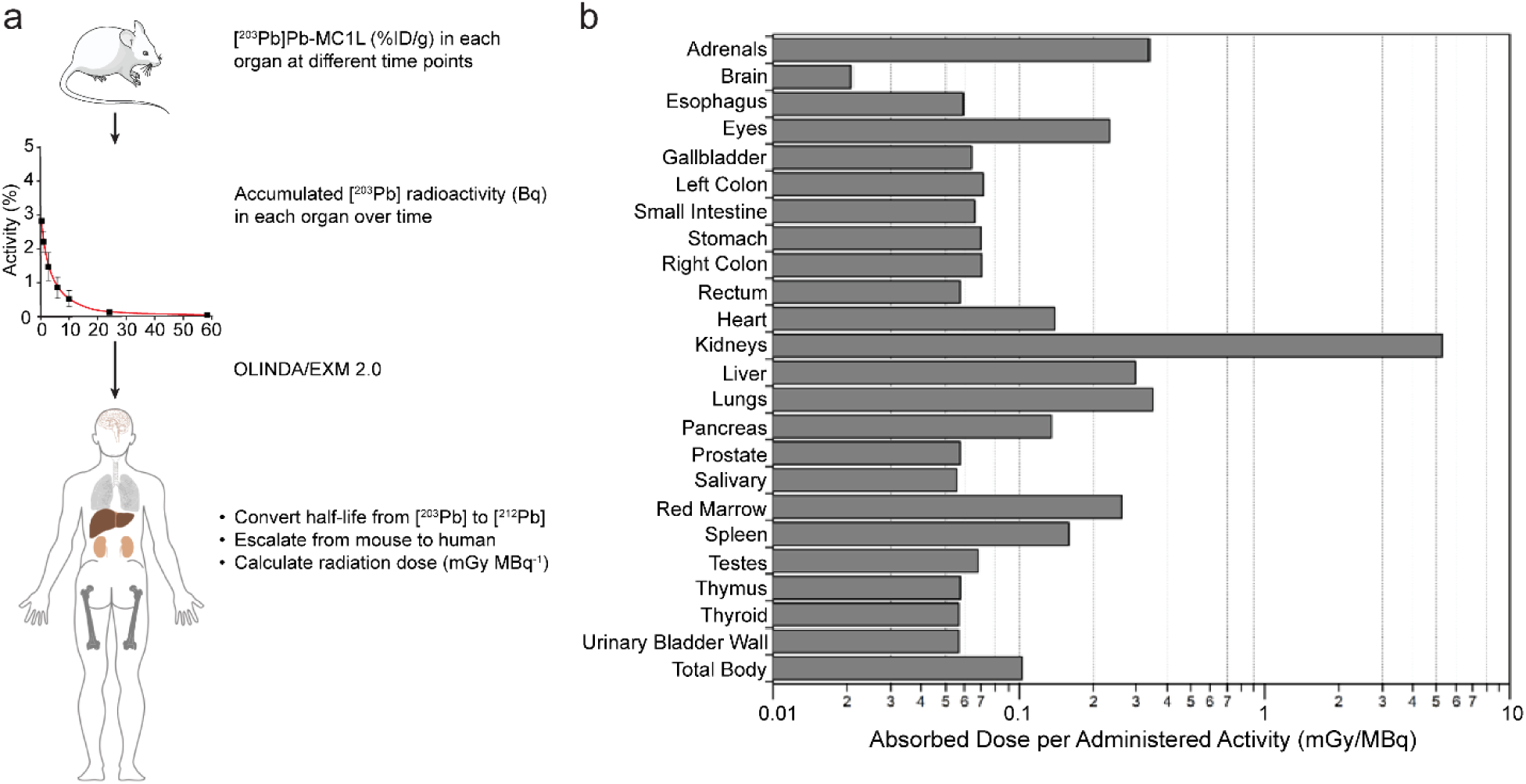
(a) Representative figure of how radiation dose was estimated in humans. (b) Dosimetry analysis of [^212^Pb]Pb-MC1L in female and male human subject Phantoms in OLINDA/EXM 2.0.

### Systemic, Cardiac, and Liver Toxicity

To identify [^212^Pb]Pb-MC1L chronic toxicity complications, the study was completed at 28 weeks after treatment, as this represents one-third of the expected lifespan of these animals (**Figure 4a**). No acute weight loss was observed in any [^212^Pb]Pb-MC1L-treated mice. However, mice treated with 6.7 MBq of [^212^Pb]Pb-MC1L demonstrated a lack of weight gain that initiated about 50 days post-treatment and was significantly different compared to the control group after 28 weeks (45.8±5.8 g vs. 59.8±11.2 g, p=0.0056, **Figure 4b**).

**Fig 4.**
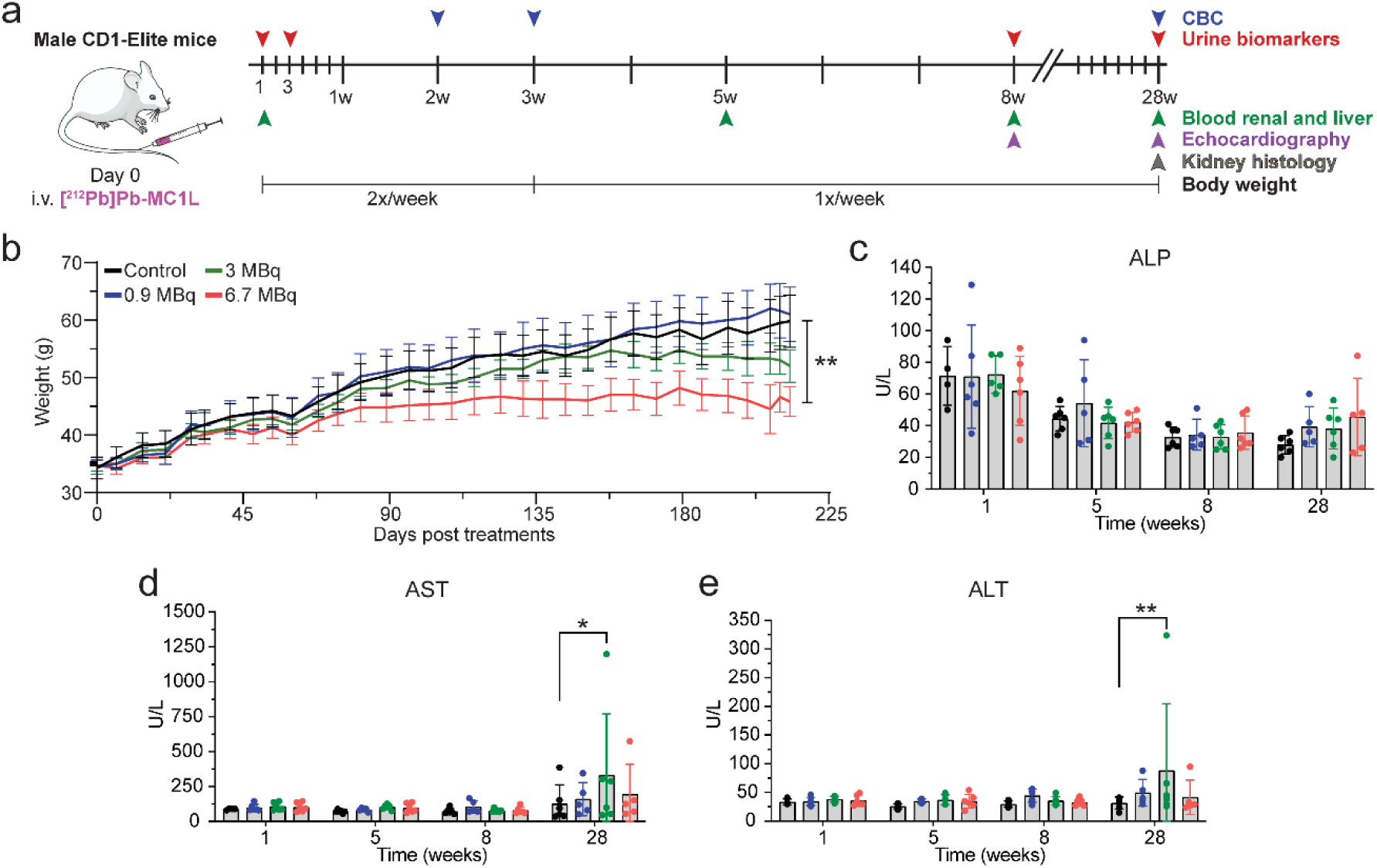
(a) Experiment design. (b) Body weight in male CD-1 Elite. (c-e) Liver function measurement by serum chemistry analysis. Statistical analysis was made by 2-way ANOVA analysis with * p<0.05.

Cardiac structure and function were analyzed by echocardiogram at 8- and 28 weeks post-treatment. No significant cardiac structural or functional changes were identified with the evaluation of heart rate, left ventricular mass, end-diastolic volume, and ejection fraction in [^212^Pb]Pb-MC1L-treated mice (**Supplemental Figure 1**).

There was no significant change in the liver injury markers ALT, AST, and ALP (**Figure 4c-e**) in the first 8 weeks post-treatment. There was a higher concentration of AST and ALT in animals treated with 3.0 MBq [^212^Pb]Pb-MC1L but not in animals treated with 6.7 MBq at 28 weeks post-treatment. However, there was no dose-dependent change in liver injury markers concentrations with levels lower than the threshold of severe hepatocellular injury based on Hy’s Law (≥ 3-fold rise in AST, ALT), which correlates with the low [^203^Pb]Pb-MC1L liver uptake observed (**Figure 2a**),

### Renal Toxicity Evaluation

Presentation of radiation-induced nephrotoxicity is usually late in both pre-clinical models and humans. Consistent with kidneys being frequently a treatment-limiting organ for RLTs, we observed rapid and high kidney uptake in our biodistribution and dosimetry studies (**Figure 2a**), which led us to perform an in-depth kidney toxicity evaluation. In the first 8-weeks post-treatment, there was no evidence of kidney toxicity by measuring the serum endogenous kidney function biomarkers BUN and creatinine. At 28 weeks post-treatment, BUN and sCr were significantly elevated in animals treated with 3.0 and 6.7 MBq, but not 0.9 MBq [^212^Pb]Pb-MC1L compared to control mice (**Figure 5a-b**). To address the low sCr sensitivity for kidney dysfunction detection in mice[19], we also measured serum cystatin C as a superior surrogate marker of kidney function at 28 weeks. Elevated cystatin C was only observed in animals injected with 6.7 MBq [^212^Pb]Pb-MC1L, but not in other groups (**Figure 5c**). Kidney histological analysis at 28 weeks following [^212^Pb]Pb-MC1L treatment demonstrated a significant increase in tubular injury, interstitial inflammation, glomerular changes, and fibrosis in mice treated with 6.7 MBq compared to control mice (**Figure 5d-e**). There was some evidence of tubulointerstitial injury in other treated groups that appeared to be dose-dependent; however, it was non-statistically significant (**Figure 5e**). Using an injury score from 0-4 in all histological changes, the injury to the glomerular region was less severe than to the tubular and interstitial regions (**Figure 5e**). Our findings highlight that kidney toxicity is more severe in the tubulointerstitium, is dose-dependent, and presents late after treatment. Moreover, we demonstrated that current diagnostic tools to evaluate nephrotoxicity are unchanged early after treatment and present late once significant irreversible kidney damage and fibrosis occur. Using our findings and based on exponential-power regression similar to Barone et al.[20], we found that 50% normal tissue complication probability (NTCP) for our study is approximately 8.0–9.5 Gy, depending on the biological data used for modeling (fibrosis, late creatinine rise, late uNGAL rise). These results are approximately consistent with the range of expected RBE factors, and our results may inform subsequent murine experimental designs for alpha-emitting radiopharmaceuticals.

**Fig 5.**
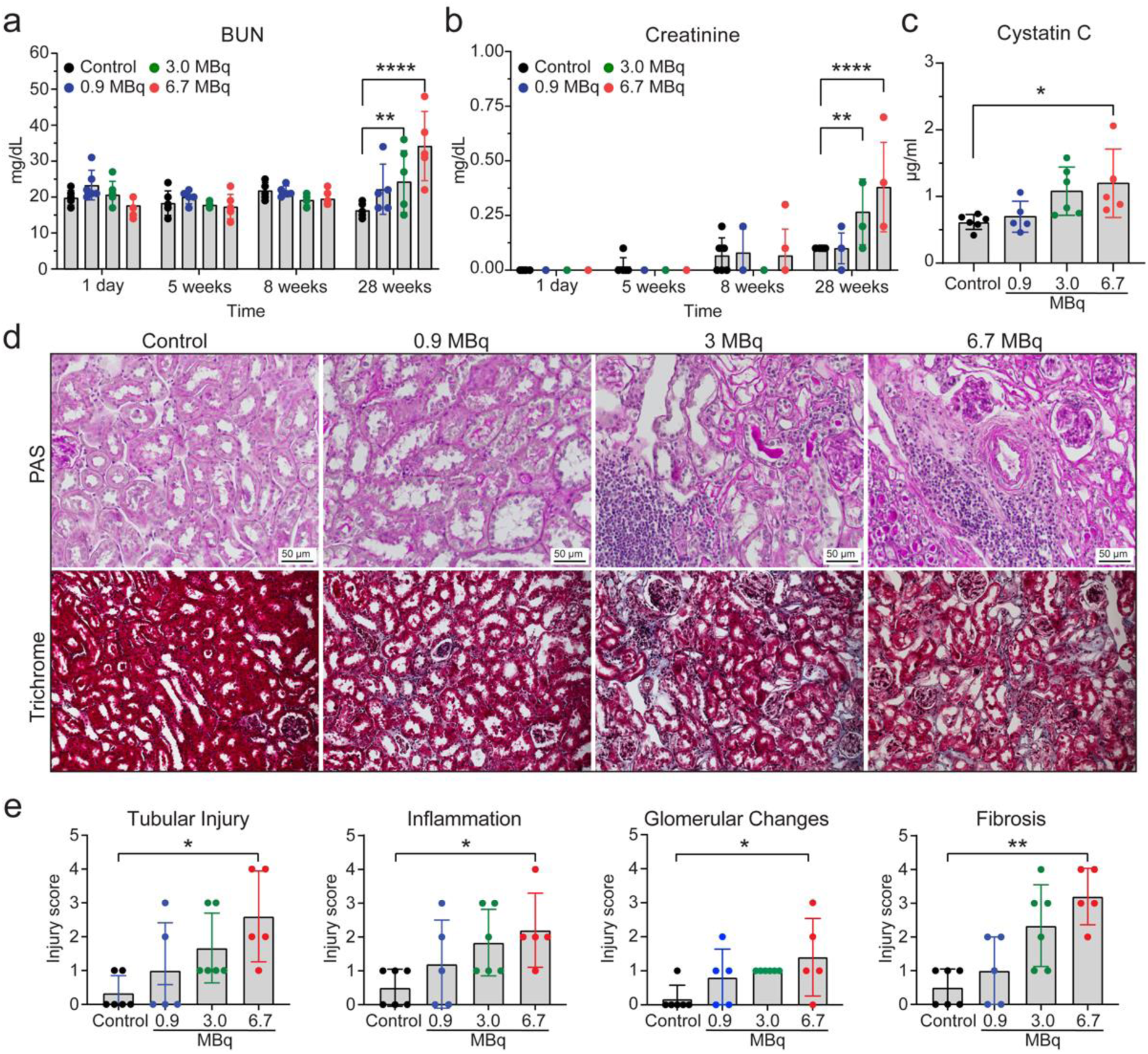
(a) BUN (Blood Urea Nitrogen), (b) Creatinine, and (c) Cystatin C kidney function markers. (d) Representative images of kidney sections stained by PAS (Periodic acid-Schiff) and Trichrome, scale bar 50 μm. (e) Tubular Injury (TI), Interstitial Inflammation (IF), Glomerulosclerosis (GS), and Fibrosis (FIB) histological scoring of kidney sections from CD-1 Elite mice at the end-of-study. Statistical analysis was made by 2-way ANOVA analysis for serum biomarkers and Kruskal-Wallis test for histological score analysis, with * p<0.05.

### Urine Biomarkers for AKI and Correlation Analysis

To test the hypothesis that RLTs cause early tubular injury that is not identified by conventional kidney function markers and that the severity of the earlier injury could correlate with long-term kidney toxicity, we measured a panel of novel biomarkers of tubular injury (NGAL and KIM-1) and tubular health (EGF) in [^212^Pb]Pb-MC1L treated mice. We collected urine in the acute injury phase (days 1 and 3) and prospectively at 8-and 28 weeks after [^212^Pb]Pb-MC1L injection to measure urine NGAL, KIM-1, and EGF levels (**Figure 6**). We also measured urine protein to creatinine ratio (UPC), a maker of hyperfiltration and a surrogate marker of glomerular damage routinely used to identify CKD progression risk[21].

**Fig 6.**
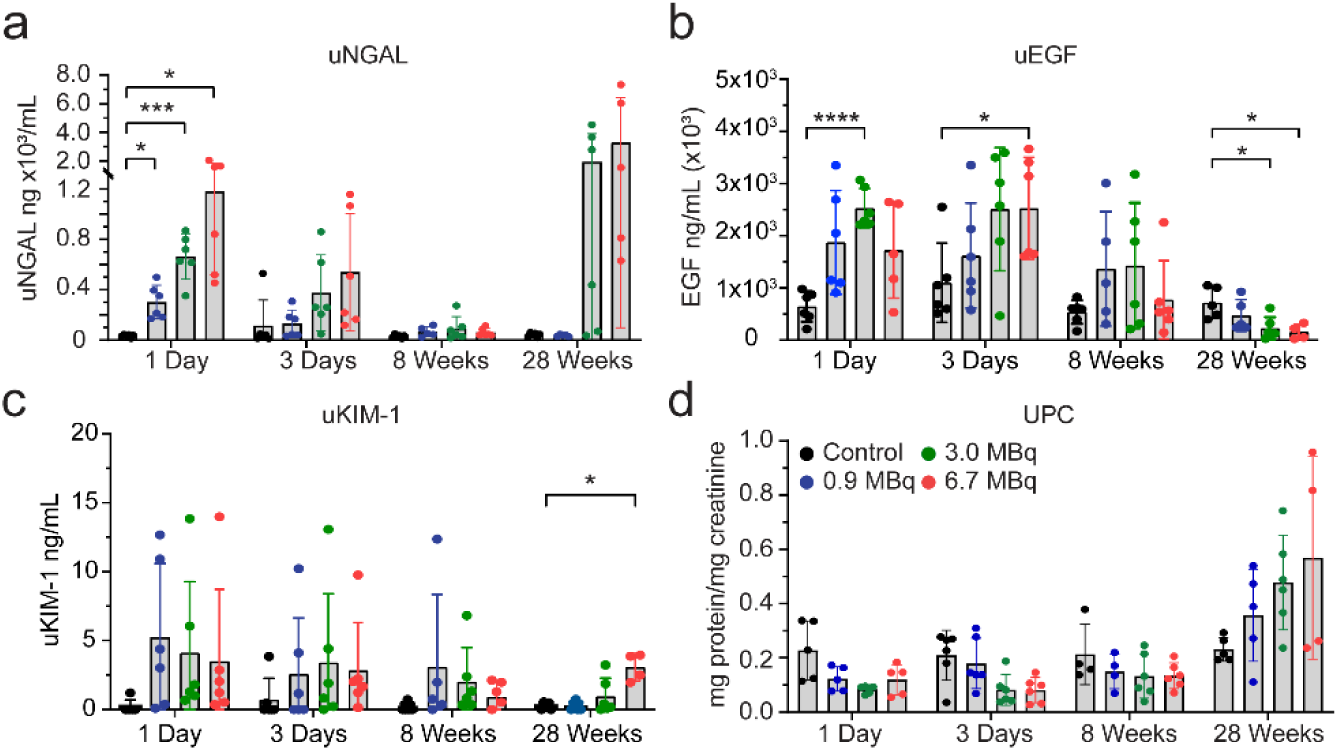
Urine biomarkers (a) NGAL (Neutrophil gelatinase-associated lipocalin. (b) EGF (Epidermal Growth Factor). (c) KIM-1 (Kidney Injury molecule 1). (d) UPC (urine protein: creatinine ratio). Statistical analysis was made by two-way ANOVA mixed-effect analysis with * p<0.05.

There was a dose-dependent significant increase in urine NGAL (uNGAL) excretion on day 1 compared to the control group (**Figure 6a**). The increase in uNGAL excretion in mice with unchanged kidney function biomarkers (BUN, sCr, Cystatin C) or proteinuria suggests the presence of subclinical AKI caused by tubular injury. Moreover, the early increase in uNGAL positively correlated with the injected [^212^Pb]Pb-MC1L dose (r=0.89, p=<0.0001), kidney absorbed dose (r=0.89, p=<0.001) and with the late histological alterations of TI (r=0.72, p=0.0002), IF (r=0.66, p=0.001), GC (r=0.58, p=0.004) and FIB (r=0.75, p=<0.0001) 28 weeks post-treatment (**Figure 7, yellow rectangle, Supplemental Figure 2**). The correlation between uNGAL secretion and late kidney findings could suggest that the severity of tubular cell injury could be a driver of CKD progression.

**Fig 7.**
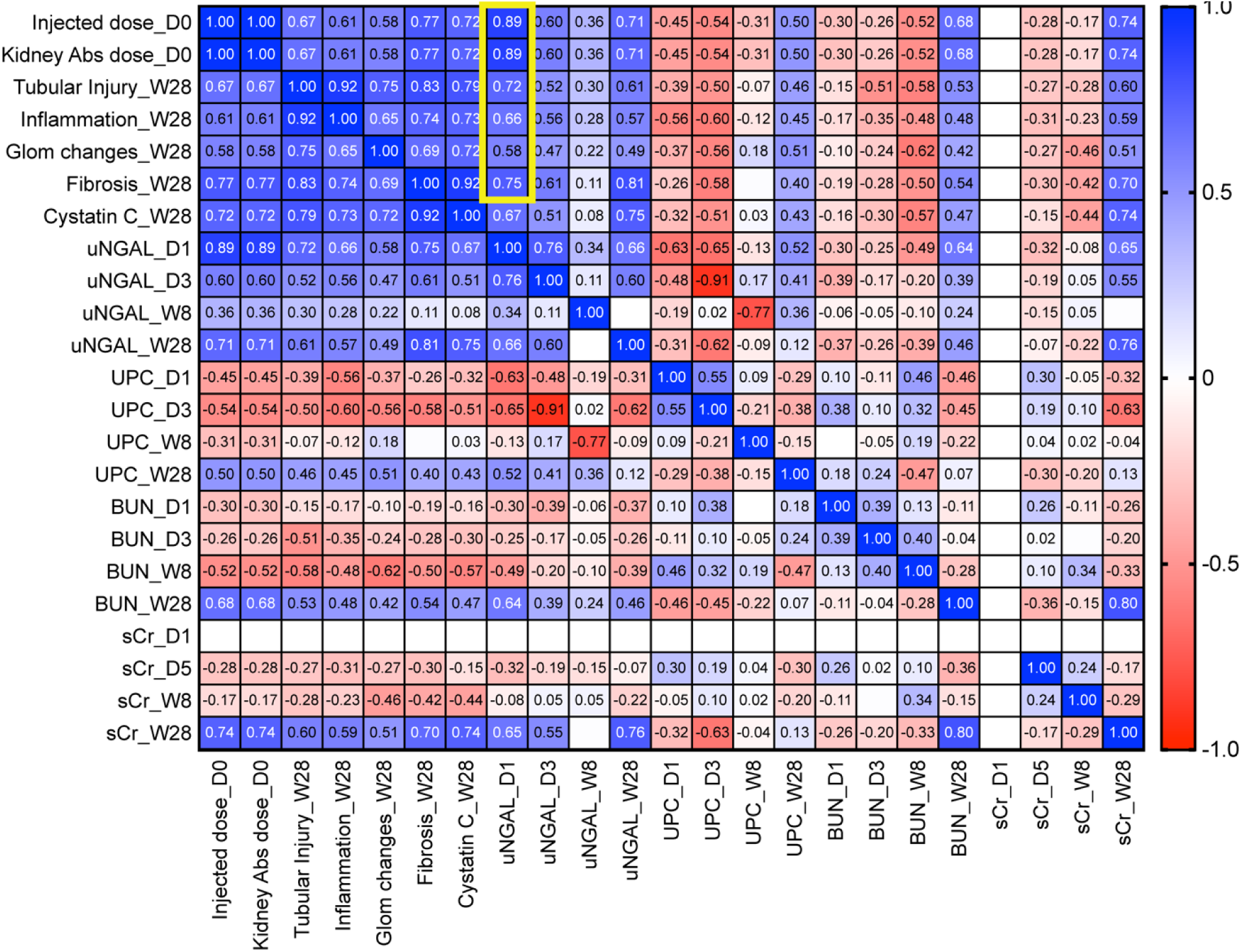
Correlation analysis of injected dose, kidney damage scores, uNGAL (yellow rectangle), UPC, BUN, and sCr at each time point. Statistical analysis was made by computed nonparametric Spearman correlation for urine biomarkers and multiparameter correlation.

High urinary levels of EGF (uEGF) reflect functional tubular mass and regeneration potential[22] and promote tubular cell proliferation modulating the recovery from acute kidney injury[23]. There was a significant increase in uEGF secretion on days 1 and 3 post-treatment in mice receiving 3.0 MBq and 6.7 MBq, respectively. Conversely, there was a lower urinary secretion at 28 weeks in mice treated with 3.0 and 6.7 MBq, consistent with tubulointerstitial damage (**Figure 6b**), suggesting lower tubular health[22]. The proximal tubule injury biomarker KIM-1 appeared to be elevated in acute and chronic stages. However, the concentrations were highly variable; the urinary excretion did not have an apparent dose-dependent increment in the acute phase and was only significant at 28 weeks in the group receiving the highest dose (6.7 MBq) (**Figure 6c**).

UPC ratio was not statistically significant at any time point. However, it showed a dose-dependent trend toward increasing proteinuria at 28 weeks when there was already evidence of kidney dysfunction by sCr and cystatin C (**Figure 6d**). Our results demonstrated that biomarkers of tubular injury and tubular health are more sensitive than endogenous biomarkers of kidney function to detect early [^212^Pb]Pb-MC1L mediated injury.

### Bone Marrow Toxicity

To evaluate bone marrow toxicity, we performed CBC analysis at 3- and 28 weeks following the injection of [^212^Pb]Pb-MC1L in male CD-1 Elite mice (n=6) (**Figure 4a**). At 3 weeks post-injection, there was no significant change in any indicative biomarker of acute bone marrow toxicity (*i.e.,* WBC, neutrophil, lymphocyte, hemoglobin, and RBC), indicating no acute hematotoxicity resulted from [^212^Pb]Pb-MC1L treatment (**Figure 8**). At the conclusion of the study (28 weeks), no significant change was observed in WBC, neutrophils, or lymphocytes in any group of mice treated with [^212^Pb]Pb-MC1L compared with the control cohort (**Figure 8a-c**). However, there was an isolated reduction of hemoglobin and RBC at 28 weeks post-injection of [^212^Pb]Pb-MC1L in mice treated with 3 and 6 MBq (**Figure 8d-e**). Based on renal findings with evidence of CKD by histology and functional biomarkers at this stage, the isolated changes in hemoglobin and RBC could be secondary to anemia of CKD and not due to bone marrow toxicity.

**Fig 8.**
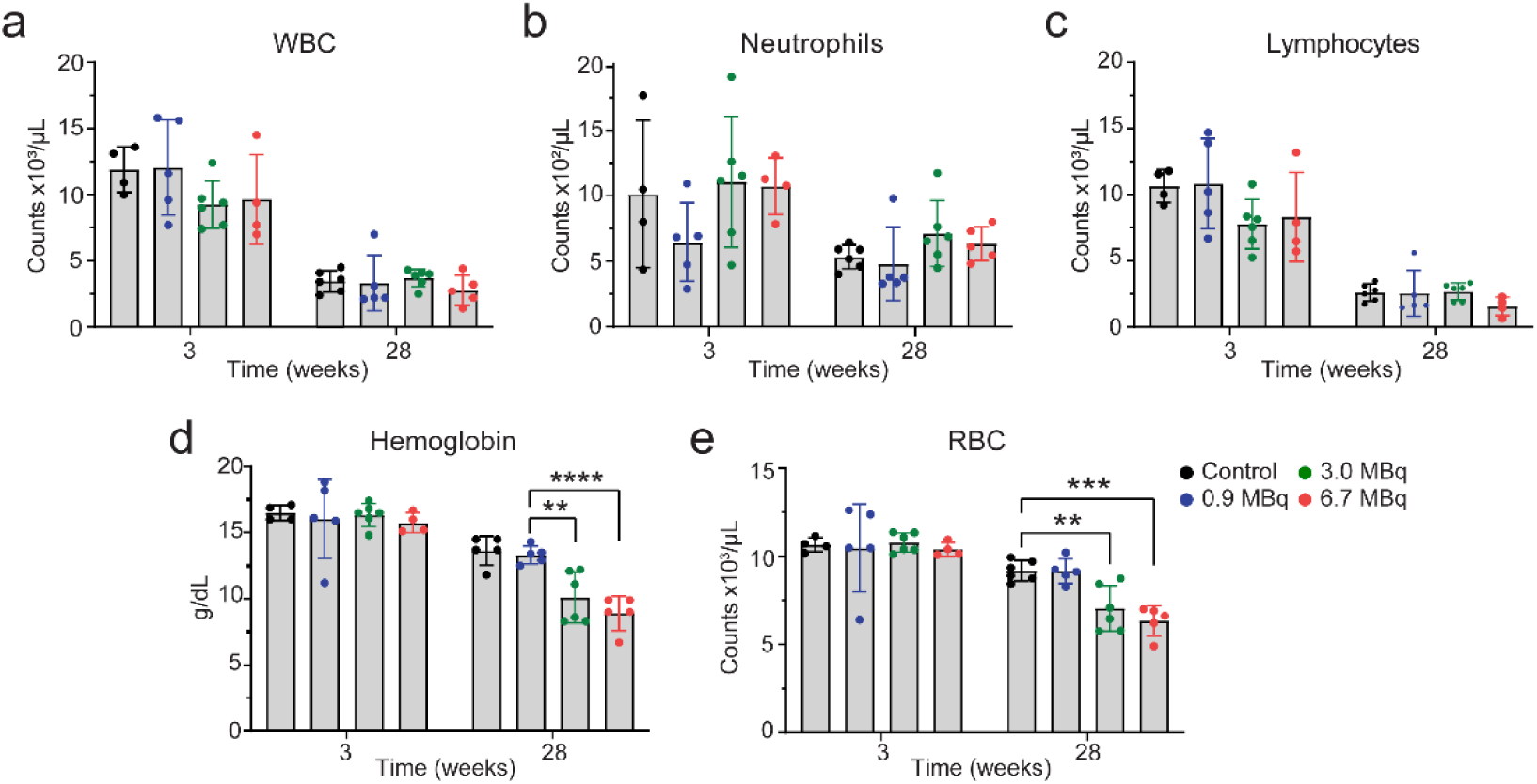
Complete blood counts (a) white blood cells (WBC). (b) Neutrophils. (c) Lymphocytes. (d) Hemoglobin. (e) Red blood cells (RBC). Statistical analysis was made by 2-way ANOVA analysis with * p<0.05.

## Discussion

In this study, we report for the first time the use of urinary biomarkers of tubulointerstitial disease to identify subclinical AKI and predict CKD associated with α-emitter RLT in a murine model. Using [^203^Pb]Pb-MC1L to characterize the distribution and absorbed dose of [^212^Pb]Pb-MC1L, kidneys appear to have the highest uptake. The dose deposition in kidneys from injection of 0.9, 3.0, and 6.7 MBq [^212^Pb]Pb-MC1L were, on average, 2.8, 8.9, and 20 Gy, respectively. The injected activities were selected to cover low, medium, and high kidney doses intended to induce AKI. A dose escalation toxicity study in mice was well tolerated based on body weight measurement with lack of significant hematotoxicity and liver toxicity. However, we observed the development of late CKD in mice treated with higher [^212^Pb]Pb-MC1L doses based on the profile change of kidney function biomarkers and the kidney histological alterations found 28 weeks after treatment. There was a significant dose-dependent urinary excretion of the tubular injury biomarker uNGAL the day after [^212^Pb]Pb-MC1L treatment that highly correlated with the severity of tubulointerstitial injury and late kidney function biomarkers. Despite the lack of early changes in conventional endogenous kidney function biomarkers (BUN, sCr, and serum cystatin C), the early concurrent increase in tubular injury suggests that early sub-clinical tubular injury could explain the long-term renal dysfunction observed at 28 weeks.

Dosimetry is a strategy that facilitates the determination of doses administered to normal organs, including those more sensitive to RLT toxicity. In the present study, we administered [^212^Pb]Pb-labeled peptide [^212^Pb]Pb-MC1L via *i.v.* injection to deliver α-radiation. MC1L is a synthetic peptide of 7 amino acids cyclized via Cu-mediated click chemistry. As an analog of alpha-melanocortin stimulating hormone (α-MSH), MC1L binds with melanocortin receptor 1 (MC1R) overexpressed in metastatic melanoma. Within this paradigm, the 10.6-hour half-life of [^212^Pb] decay matches the relatively shorter biological half-life of small peptides compared with the long biological half-lives of antibodies[24]. Cyclotron-produced gamma-ray-emitting radionuclide [^203^Pb] was used as an elementally identical imaging surrogate for [^212^Pb][7]. [^203^Pb]Pb-MC1L surrogate was injected in the same animal model to determine the *in vivo* biodistribution for calculating the dosimetry of [^203^Pb]Pb-MC1L in normal organs. Our prior studies have demonstrated excellent chelation kinetics and stability with PSC for [^212^Pb] and [^212^Bi][8]. However previous literature has shown that during a fraction of [^212^Pb] disintegrations, the daughter [^212^Bi] cations are released from the chelator[25]. Based on the known kinetics of bismuth, it is likely that some released ions will accumulate and decay in liver and kidneys. Therefore, doses to these structures may be underestimated when utilizing [^203^Pb] as a dosimetric surrogate without accounting for these effects.

Due to its rapid clearance, we observed low [^203^Pb]Pb-MC1L liver and blood uptake and retention. We did not observe significant liver toxicity after [^212^Pb]Pb-MC1L treatment like in previous reports using [^177^Lu]Lu-DOTATATE[26]. Some cases of hematological toxicity (grade 3/4) and dose-dependent acute hematological toxicity have been reported in patients receiving [^177^Lu]Lu-DOTATATE[27] and [^225^Ac]Ac-DOTATOC[28] therapy. Hematological toxicity after RLTs increases in patients with baseline nephrotoxicity (transient or persistent elevation in sCr), probably due to the prolonged circulation time of the radiopeptide in patients with poor kidney function, which increases bone marrow exposure to radiation[29]. Our study showed a late reduction in hemoglobin and RBC 28 weeks after [^212^Pb]Pb-MC1L treatment in mice with evidence of kidney dysfunction based on BUN and sCr measurements. Because anemia is common in patients with CKD due to reduced kidney erythropoietin production, our findings suggest the late reduction in hemoglobin and RBC after [^212^Pb]Pb-MC1L treatment was secondary to anemia of CKD.

The understanding of RLTs-associated nephrotoxicity in patients is limited and primarily evaluated by measuring endogenous kidney functional biomarkers, like sCr, that do not assess the severity or localization of tubulointerstitial damage. During steady-state conditions, sCr is commonly used to estimate GFR. More than half of the kidney mass must be impaired to detect an increase in sCr as a kidney dysfunction biomarker[30], explaining the late increase in functional biomarkers after [^212^Pb]Pb-MC1L treatment as evidence of the histological CKD progression. Moreover, sCr levels are affected by tubular secretion and reabsorption, kidney generation, extrarenal elimination, certain medications, and patient gender, age, diet, and muscle mass, which results in an overestimation of GFR and does not reflect accurately intrinsic kidney injury[31]. The reports of kidney toxicity in α-emitters are limited, and sCr restrictions may explain the controversy in nephrotoxicity development. In a study of 39 patients receiving [^225^Ac]Ac-DOTATOC α-RLT, only 22 were suitable to evaluate long-term kidney function due to tumor progression or change in therapy. Over a median of 57 months (range 18-90 months), they observed an average eGFR-loss of 8.4ml/min (9.9%) per year, and treatment-related kidney failure occurred in 2 patients after a delay of >4 years[32]. The lack of accuracy in nephrotoxicity detection, mainly by sCr-based eGFR, led to the conclusion that kidney failure is independent of administered radioactivity and other clinical risk factors are significant or the main contributors[32–34]. Following findings in humans, we did not observe any early changes in the conventional kidney function biomarkers BUN and creatinine. We only found late significant changes at 28 weeks in mice treated with [^212^Pb]Pb-MC1L at 3.0 and 6.7 MBq, consistent with a kidney-absorbed dose of ∼8.9 and ∼20 Gy, respectively. Even though no RBE scaling was used in this work to avoid ambiguity, and all doses are presented as physical absorbed dose to the structure of interest, these results are in accordance with pre-clinical studies using [^213^Bi]Bi-DOTATATE (RBE of 4–5 for acute renal toxicity), where the calculated renal LD_50_, at which 50% of the treated animals would develop acute nephrotoxicity, was 20±8 Gy[35]. Kidney functional biomarkers (BUN and sCr) do not directly detect injury in the tubular epithelium, the leading site of injury in most forms of AKI, that, without treatment, can progress to CKD[13]. In line with radiopeptide reabsorption by proximal tubular cells, pre-clinical data in mice and rat models have shown evidence of long-term tubulointerstitial disease after β-RLTs and α-RLT in the absence of severe glomerular pathology or sCr changes[11,12]. In this context, the delayed change in kidney function biomarkers and the lack of information on tubular injury results in poor accuracy in diagnosing acute structural kidney damage associated with RLT treatment and could result in missing valuable treatment windows to prevent CKD progression.

Biomarkers of tubular injury and health[36] are more frequently used to diagnose AKI before the loss of filtration function (kidney function biomarkers changes), supporting their potential incorporation into nephrotoxic evaluation in RLTs patients. Urinary biomarkers of tubular health (EGF) and injury (KIM-1 and NGAL) measurements in mice treated with [^212^Pb]Pb-MC1L showed an early increase in NGAL and EGF excretion. NGAL is a protein reabsorbed by proximal tubules and released by the damaged distal tubules in response to acute tubular injury, which can be detected in the urine within hours of tubular injury[37]. Thus, an increase in urine NGAL reflects both distal tubular injury that causes an increase in NGAL production to the urine and systemic circulation and proximal tubular damage that results in a lack of NGAL reabsorption by proximal tubular cells[38]. We demonstrate that uNGAL excretion was significantly increased after [^212^Pb]Pb-MC1L treatment with a dose-dependent pattern on day 1 after treatment. Moreover, its increased urinary excretion was highly correlated with tissue damage at 28 weeks, similar to the long-term serum NGAL increase observed in mice receiving the α-emitter [^213^Bi]Bi-DOTATATE[35]. The early elevation in uNGAL indicates a dose-dependent [^212^Pb]Pb-MC1L-induced tubular injury that correlates with long-term kidney toxicity. The observed early increase in tubular injury biomarkers with a lack of changes in kidney function biomarkers is used to diagnose a subclinical AKI condition[39]. This condition is usually asymptomatic and has been associated with a 2-3-fold increased risk of death or the need for kidney support therapy compared to patients with normal levels of sCr and tubular damage biomarkers[40]. EGF expression is specific for the ascending limb of Henle and distal tubules and has been associated with enhanced tubular cell regeneration and repair, accelerating kidney recovery after injury[23,41]. uEGF is decreased in various kidney diseases in humans, including patients with CKD[42]. We found a significant initial increase in uEGF excretion in the 3 MBq and 6.7 MBq that was then significantly reduced at 28 weeks after treatment suggesting an adequate initial response to injury with a lower late tubular reserve 28 weeks after treatment compared to the control group. Finally, KIM-1 is a protein expressed in proximal tubular epithelial cells rarely expressed in organs other than kidneys, whose increased urine level is a sensitive indicator for early proximal tubular injury, particularly for ischemic or nephrotoxic AKI patients[43]. We found a trend towards an increase in uKIM-1 after [^212^Pb]Pb-MC1L treatment that persisted and was significant in the mice treated with 6.7 MBq after 28 weeks. The urinary biomarkers patterns suggest that proximal tubule cells might not be the only tubular segment affected by [^212^Pb]Pb-MC1L treatment.

We recognize some limitations in our study design. Studies evaluating long-term radiation-induced toxicity can be extended up to 36-week post therapy[44], and our study concluded at 28 weeks (∼7 months). However, the average mouse lifespan is about 24 months[45], and a follow-up of 7 months post-treatment represents approximately one-third of their lifespan. In murine CKD models, where it is expected to have a gradual renal function decline, mice are followed between 1-6 months post-injury to evaluate chronic changes in tissue, including fibrosis[46]. Because mice treated with 6.7 MBq had significant changes in kidney function biomarkers suggesting clinically meaningful renal function decline, we concluded our study at 28 weeks to ensure adequate survival in each group to compare histological findings. We found significant fibrosis in that group (score 3 of 4), supporting adequate time to identify fibrosis in this model.

The results from this dose-escalation pre-clinical toxicity study propose that increased uNGAL secretion could be an additional diagnostic tool to identify patients developing α-RLT subclinical AKI and at risk of CKD. The changes in uEGF suggest that proximal tubule cells might not be the only tubular segment affected by α-RLT and highlight how potential early tubular damage could be an essential driver of long-term α-RLT-related nephrotoxicity. Together with the conventional clinical predictors of kidney health (sCr, cystatin C, UPC), urine biomarkers of tubular injury (i.e., NGAL) and tubular health (i.e., EGF) in combination with dosimetry, could be used to personalize α-RLT and provide higher doses to those patients without evidence of tubulointerstitial injury. On the contrary, in patients where therapy is still required and highly beneficial, it would empower practitioners to discuss the long-term risk with patients and ensure early and close CKD follow-up to improve their outcomes, as previously proposed. More studies using dosimetry and evaluating biomarkers of tubulointerstitial disease in RLTs are needed to identify timing of tubular injury at specific renal dosages and guide future toxicity mechanistic studies. This approach could facilitate tumor treatment maximization, development of nephroprotective therapies, and identification of patients at risk of long-term toxicity.

## Statements & Declarations

### Author Contributions

ML, CR-P: data analysis and interpretation; writing-original draft. DLiu: experiment design, sample collection, and processing. SAG: mouse and human dosimetry analysis and interpretation. GV-M, GM-A: kidney biomarkers measurement. DLee: provided consultation for Perspective Therapeutics. PR: blind PAS histological analysis. BMM: sample collection and processing. EAS: Radionuclide generation. RMW: Cardiotoxicity analysis. SCL: blind Trichome histological analysis. FLJ: writing - review and editing. DZ-O, MKS: conceptualization, methodology, writing - review and editing, supervision.

### Data Availability

The datasets generated during and/or analyzed during the current study are available from the corresponding author upon reasonable request.

### Ethics approval

All animal studies followed the Guide for the Care and Use of Laboratory Animals and were approved by the University of Iowa Animal Care and Use Committee.

## Footnotes

1 Low-energy beta emitters ([^177^Lu], [^67^Cu], [^131^I]), does not include higher energy emitters ([^90^Y], [^89^Sr], [^32^P], etc.)

