## Supplemental Material and Figures for "Pre-clinical Evaluation of Biomarkers for Early Detection of Nephrotoxicity Following Alpha-particle Radioligand Therapy"

**European Journal of Nuclear Medicine and Molecular Imaging**

Mengshi Li,[1*] Claudia Robles-Planells,[2*] Dijie Liu,[1] Stephen A. Graves,[3] Gabriela Vasquez-Martinez,[2] Gabriel Mayoral-Andrade,[2] Dongyoul Lee,[4] Prerna Rastogi,[5] Brenna M. Marks,[1] Edwin A. Sagastume,[1] Robert M. Weiss,[6] Sarah C. Linn-Peirano,[2,7] Frances L. Johnson,[1] Michael K. Schultz,[1,3,8#] Diana Zepeda-Orozco[2,9,10#]

[1]Viewpoint Molecular Targeting, Inc., Coralville, IA, USA

[2]Kidney and Urinary Tract Center, Abigail Wexner Research Institute at Nationwide Children's, Columbus, Ohio, USA

[3]Department of Radiology, The University of Iowa, Iowa City, IA, USA.

[4]Department of Physics and Chemistry, Korea Military Academy, Seoul, Republic of Korea

[5]Department of Pathology, The University of Iowa, Iowa City, IA, USA

[6]Department of Internal Medicine, The University of Iowa, Iowa City, IA, USA

[7]Department of Veterinary Biosciences, The Ohio State University College of Veterinary Medicine Columbus, OH

[8]Department of Radiation Oncology, Free Radical, and Radiation Biology Program, The University of Iowa, Iowa City, IA, USA

[9]Department of Pediatrics, The Ohio State University College of Medicine, Columbus, Ohio, USA.

[10]Division of Nephrology and Hypertension, Nationwide Children's Hospital, Columbus, Ohio, USA.

City, IA, USA.

[*]Co-First Authors

[#]Co-Corresponsing Authors

Diana Zepeda-Orozco:

Michael K. Schultz:

### Supplemental Methods

#### Reagents and Materials

Lead-212 ($^{212}$Pb) was obtained *via* a $^{224}$Ra/$^{212}$Pb generator from the National Isotope Development Center/Oak Ridge National Laboratory (Oak Ridge, TN). Lead-203 ($^{203}$Pb) was provided by Lantheus Medical Imaging (North Billerica, MA). Male CD1-Elite mice were obtained from Charles River Laboratories (Wilmington, MA). Pb resin (a chromatographic resin used for Pb isotope purification) was provided by Eichrom Technologies, Inc. (Lisle, IL). A National Institute of Standards and Technology (NIST) traceable $^{232}$U/$^{212}$Pb source in a hermetically sealed ampoule was obtained from Eckert & Ziegler (Cat# 7432, Eckert & Ziegler, Valencia, CA, USA). Other reagents and chemicals were purchased from ThermoFisher Scientific (Waltham, MA).

#### Radiosynthesis of [$^{203}$Pb]Pb-MC1L and [$^{212}$Pb]Pb-MC1L

[$^{203}$Pb]Pb-MC1L surrogate was used to determine the biodistribution of Pb-MC1L in normal organs. Radiolabeling of MC1L was conducted according to published methods[6]. $^{203}Pb^{2+}$ was purified on 50 mg Pb resin and eluted by 2 mL 0.5 M sodium acetate solution (NaOAc, pH=6) into the reaction vessel containing 20 nmole MC1L precursor and 0.5 M NaOAc (pH=4) buffer to adjust final pH to 5.4-5.5. The reaction vessel was heated at 85°C for 30 min, followed by the removal of unreacted $^{203}Pb^{2+}$ on StrataX C-18 SPE cartridge (Phenomenex, Torrance, CA). [$^{203}$Pb]Pb-MC1L final product on StrataX C-18 SPE cartridge was collected in 50% EtOH in saline. Radiolabeling of [$^{212}$Pb]Pb-MC1L was performed according to previously published methods[6]. Briefly, $^{212}Pb^{2+}$ and daughters were eluted from the $^{224}$Ra/$^{212}$Pb generator in 2 M hydrochloric acid (HCl). The $^{212}Pb^{2+}$ solution was loaded onto a pre-packed SPE column containing 50 mg Pb resin. Purified $^{212}Pb^{2+}$ was eluted off the Pb resin by 2 mL 0.5 M NaOAc buffer (pH=6) directly to the reaction vessel containing 26 nmole MC1L, 2 mg sodium ascorbate, 100 µL 200-proof ethanol (EtOH), and 290 µL 0.5 M NaOAc buffer (pH=4). Following the reaction at 80°C for 30 min, purification of [$^{212}$Pb]Pb-MC1L (*i.e.*, removal of any unreacted $^{212}Pb^{2+}$) was conducted on StrataX C-18 SPE cartridge. Purified [$^{212}$Pb]-MC1L was then collected using 1 mL 50% EtOH in saline. Radioactivity of [$^{212}$Pb]Pb-MC1L sample was measured on a sodium iodide (NaI) gamma spectrometer (ORTEC, Oak Ridge, TN) and ionization chamber (IC) dose calibrator (CRC-15R,

Capintec, Florham Park, NJ) that were calibrated with NIST-traceable $^{232}U/^{212}Pb$ standard source as previously described[8].

**Dosimetry analysis of [$^{212}$Pb]-MC1L using [$^{203}$Pb]-MC1L data**

Using the biodistribution data of [$^{203}$Pb]Pb-MC1L surrogate, the absorbed dose from injection of [$^{212}$Pb]Pb-MC1L in mouse and human were calculated using OLINDA/EXM 2.2. Mass correction in OLINDA was utilized based on the average organ masses for animals within the ex vivo biodistribution studies.

Murine organ biodistribution data for [$^{203}$Pb]-MC1L are summarized in **Supplemental Table 2**. From these data percent administered activity per organ (%AA/organ) for $^{212}$Pb-MC1L was calculated from the percent injected dose per gram (%ID/g) corrected for the decay of [$^{203}$Pb] as follows:

$$\%AA/organ = (\%ID/g) * (m_{organ}) * (e^{-\lambda t}) \qquad \textit{Eq. 1}$$

where $m_{organ}$ is the average organ mass, and $e^{-\lambda t}$ is the Lead-212 decay factor for the data collection time-point. Resulting %AA/organ values are summarized in **Supplemental Table 3**.

To determine time-integrated activity coefficients for each organ, two methods were utilized.

*Method 1*: Each organ time activity curve (%AA/organ vs. time) was fit using bi-exponential regression. Time activity curves and associated bi-exponential fits are shown in **Supplemental Figure 3.** The resulting fit parameters $A_1, \lambda_1, A_2$, and $\lambda_2$ describe the initial distribution and clearance, followed by a much slower excretion phase. These fit parameters are summarized in **Supplemental Table 4.**

*Method 2*: Trapezoidal integration was used for time-integrated activity coefficient determination by assuming constant uptake between t=0 and t=0.5 h, and physical decay following the last time-point.

The average of time-integrated activity coefficients from methods 1 and 2 was utilized for all subsequent dose calculations. Resulting time-integrated activity coefficients are summarized in **Supplementary Table 4**.

Murine organ absorbed dose was determined using OLINDA v2.2 from the determined time-integrated activity coefficients. The 35-gram mouse model was utilized for s-value lookup, and organ masses were scaled per the measured organ masses in **Supplemental Table 2**. Notably, decay chain calculations are not currently supported in OLINDA 2.2 for [$^{212}$Pb], so separate calculations were performed for the four relevant radioisotopes ([$^{212}$Pb], [$^{212}$Bi], [$^{212}$Po], [$^{208}$Tl]) with appropriate consideration of branching ratios, and the results were summed for final dose specification. Blood, adrenals, muscle, skin, and eyes are not implemented as source/target tissues in the OLINDA 2.2 murine models, and therefore dose to these tissues was calculated assuming local alpha particle energy deposition, with an average alpha energy of 7.8063 MeV per [$^{212}$Pb] decay. Beta and gamma self-dose contributions were not calculated for these tissues.

Based on the assumption that the early and late exponential clearance phases in the kidneys describe residence in the glomeruli and tubules, respectively, it indicates that renal activity residence is approximately 9.9% glomerular and 90.1% tubular. The sub-organ s-values presented by Hobbs et al. can be used to estimate the associated tubular and glomerular dosimetry of [$^{212}$Pb]Pb-MC1L in mice. Correcting for kidney mass differences between our population of mice, and those used to generate the s-values described by Hobbs et al. (~0.769 multiplicative correction factor), we estimate that the tubular dose per administered activity is 6.31 Gy/MBq, and the glomerular dose per administered activity is 16.3 Gy/MBq. Within these estimates, glomerular self-dose accounts for ~84.7% of the total glomerular dose, and tubular self-dose accounts for ~95.9% of the total tubular dose.

Prediction of human dosimetry from small animal studies requires consideration of fundamental interspecies biochemical differences, the fact that mice have larger organs relative to their total body mass, and the fact that biological excretion kinetics are faster in mice than in humans. Accounting for the larger relative organ masses and more rapid excretion in mice can be accomplished using the following formula:

$$TIAC_{human} = \varepsilon * TIAC_{mouse} * \left( \frac{\left(\frac{m_{organ}}{m_{WB}}\right)_{human}}{\left(\frac{m_{organ}}{m_{WB}}\right)_{mouse}} \right) \qquad Eq.\ 2$$

Conceptually this approach can be thought of as (1) assuming SUV-equivalence for initial uptake, (2) assuming mono-exponential clearance, and (3) scaling the organ activity retention by a factor of $\varepsilon$, which represents the ratio of mouse and human tissue effective clearance rate constants ($\lambda_{mouse}/\lambda_{human}$) for each organ.

To determine $\varepsilon$ for each organ, first, we consider the estimated biological excretion kinetics by the renal pathway for mice and humans. The whole-body rate constant $k$ for biological excretion is expected to be proportionate to the ratio of the renal blood filtration rate (volume per unit time) to total blood volume (assuming a majority of the administered activity distributes to a relatively fast-turnover compartment initially). Given a normal human glomerular filtration rate (GFR) of ~100 mL/min, a human total blood volume of ~6363 mL, a mouse glomerular filtration rate of 205 µl/min[41], and a mouse blood volume of 3.00 grams (see table 1), the ratio of biological excretion rate parameters for mice and humans can be estimated as:

$$\frac{k_{mouse}}{k_{human}} \approx \frac{\left(\frac{0.205}{3.00}\right)}{\left(\frac{100}{6363}\right)} = 4.35 \qquad \textit{Eq. 3}$$

Solving this equation for $k_{human} = k_{mouse}/4.35$, and substituting this result into the formula for $\alpha$, and noting the effective clearance rate parameter ($\lambda$) is equal to the sum of the biologic clearance parameter ($k$) and the decay constant for Pb-212 ($\lambda_{Pb212}$), we have:

$$\varepsilon = \frac{\lambda_{mouse}}{\lambda_{human}} = \frac{\lambda_{mouse}}{\lambda_{Pb212} + k_{human}} \approx \frac{\lambda_{mouse}}{\lambda_{Pb212} + k_{mouse}/4.35} = \frac{\lambda_{mouse}}{\lambda_{Pb212} + (\lambda_{mouse} - \lambda_{Pb212})/4.35} \qquad \textit{Eq. 4}$$

This result, combined with the equation for $TIAC_{human}$ can be used to determine the estimated human time-integrated coefficients for each organ. Murine $TIAC$ values, organ masses (human and murine), and resulting $TIAC_{human}$ values are summarized in **Supplementary Table 5**.

Special treatment was utilized for organs of clearance (kidneys and liver). For these tissues, it is often more appropriate to assume "percent injected activity per organ equivalence" rather than "SUV equivalence," due to uptake being more related to elimination kinetics rather than the more typical reversable tissue distribution. Mathematically, this treatment can be stated as:

$$TIAC_{human} = \frac{\left(\% \frac{IA}{organ} @\, t = 0\right)}{\lambda_{human}} = \frac{\left(\% \frac{ID}{g} @\, t = 0\right)_{mouse} \left(m_{organ}\right)_{mouse}}{\lambda_{mouse}/\varepsilon} \qquad \textit{Eq. 5}$$

Initial uptake for kidneys and liver was estimated from mono-exponential fitting to the murine kidney and liver %IA/organ curves (**Figure 2**). Initial uptake was estimated to be 2.39% for kidneys and 0.70% for liver. Given murine rate constants of 0.105 $h^{-1}$ and 0.083 $h^{-1}$ for kidney and liver, one obtains $\varepsilon$ values of 1.41 and 1.20, respectively. Therefore, the estimated human time-integrated coefficients are 0.322 MBq*h/MBq and 0.101 MBq x h/MBq for kidneys and liver, respectively.

Human dosimetry was estimated using OLINDA v2.2. The ICRP 89 adult male model was used for s-value lookup, and default organ masses were utilized (as listed in **Supplementary Table 7**).

#### Hematotoxicity and Blood Chemistry evaluation

Whole blood samples were collected in CD1-Elite treated mice at 2-, 3-, and 28-weeks post [$^{212}$Pb]Pb-MC1L administration to evaluate hematotoxicity by complete blood count (CBC) analysis. In CBC analysis, whole blood was collected by tail snip and diluted in sterile ice-cold PBS by a factor of 10. White blood cells (WBC), red blood cells (RBC), lymphocytes, neutrophils, and hemoglobin were measured on Siemens ADIVA hematology analyzer. Blood chemistry panel to evaluate kidney and liver function was performed in serum samples collected at 1-, 5-, 8-, and 28 weeks post [$^{212}$Pb]Pb-MC1L injection. Serum was collected in MiniCollect™ tubes (Greiner Bio-One, Fisher Scientific). Blood chemistry panels were analyzed by IDEXX BioAnalytics (North Grafton, MA). Alanine aminotransferase (ALT), aspartate aminotransferase (AST), and alkaline phosphatase (ALP) were used to identify hepatotoxicity. Blood urea nitrogen (BUN) and creatinine were used to evaluate renal function. Upon conclusion of the study at 28 weeks, cystatin C was analyzed as additional endogenous biomarkers of kidney function using Mouse/Rat cystatin C Quantikine ELISA kit (R&D Systems, Cat# MSCT0, Minneapolis, MN). Cardiac structure and function were assessed at 8- and 28 weeks post [$^{212}$Pb]Pb-MC1L treatment in CD-1 Elite mice (n=6).

#### Cardiac Toxicity Evaluation

Briefly, under light conscious sedation (midazolam, 0.15 mg subcutaneous injection), the chest was shaved, and pre-warmed acoustic coupling gel was applied. Images were acquired in parasternal short- and long-axis planes using a 40 MHz transducer coupled to a Vevo 2100 ultrasonograph (VisualSonics, Toronto). Endocardial and epicardial contours were traced manually at end-diastole and end-systole in both parasternal planes. Left ventricular mass, end-diastolic volume, end-systolic volume, stroke volume, and ejection fraction were calculated using the biplane area-length method. Heart rate was determined using electronic calipers applied to a pulse-wave Doppler interrogation of mitral inflow. The primary readouts for determining potential cardiotoxicity were heart rate, left ventricular mass, left ventricular end-diastolic volume, and left ventricular ejection fraction, reported as a decimal (e.g., 0.80).

**Urine Nephrotoxicity Biomarkers**

Urine biomarkers measurements included NGAL (ELISA, Bioporto, Cat# IT042, Hellerup, Dinamarca), KIM-1 (Quantikine ELISA, Cat# MKM100), and EGF (Quantikine ELISA, R&D Systems, Cat# MEG00, Minneapolis, MN) were measured by ELISA. Urine protein was measured using a BCA kit (Thermo Scientific, CAT#A53226, Rockford, IL) and was normalized to urine creatinine (QuantiChrom Creatinine assay kit, Cat# DICT-500). Measurements were performed according to the manufacturer's instructions.

**Figure captions**

**Fig Supp 1. Cardiac structure and function.** Echocardiography analysis of heart rate (HR), end-diastolic volume (EDV), left ventricular thickness (LVTN), end-systolic volume (EVS), Hear mass (Mass), volume/mass ratio (V/M), stroke volume (SV), cardiac output (CO) and ejection fraction (EF). Statistical analysis was made by Kruskal-Wallis test for histological score analysis, with * $p<0.05$.

**Fig Supp 2:** Correlation analysis of injected dose, kidney damage scores, serum, and urine biomarkers. Statistical analysis was made by computed nonparametric Spearman multiparameter correlation.

**Fig Supp 3.** [$^{212}$Pb]Pb-MC1L organ time activity curves calculated from [$^{203}$Pb]Pb-MC1L biodistribution data. Lines represent bi-exponential fits to the organ time activity curve data, with fit parameters $A1, \lambda 1, A2$, and $\lambda 2$. Regression was not possible for pancreas due to the increase in organ uptake after 1.5 hours.

**Fig Supp 4.** Mono-exponential regression for murine kidney and liver time-activity data.

**Fig Supp 1**

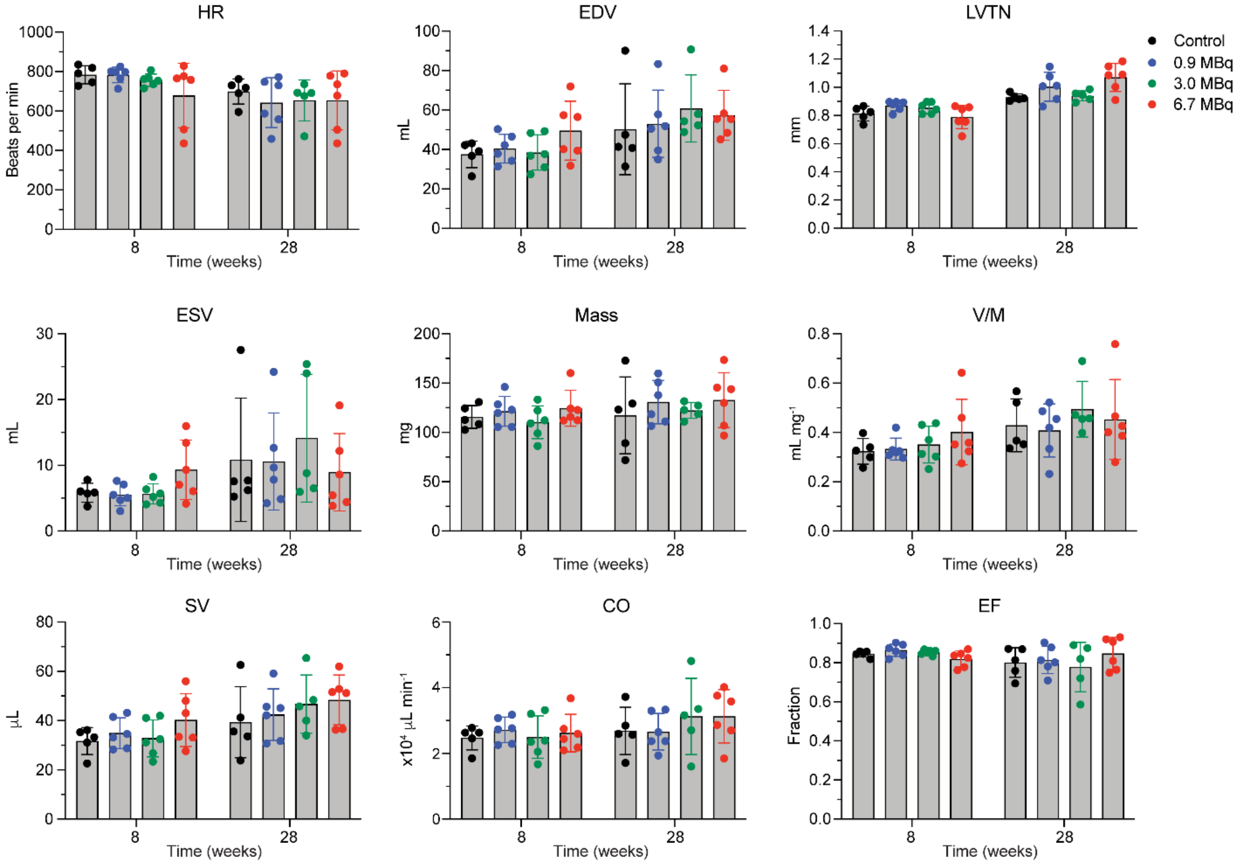
HR
Beats per min
EDV
mL
LVTN
mm
Control
0.9 MBq
3.0 MBq
6.7 MBq
ESV
mL
Mass
mg
V/M
mL mg⁻¹
SV
µL
CO
x10⁴ µL min⁻¹
EF
Fraction
Time (weeks)

**Fig Supp 2**

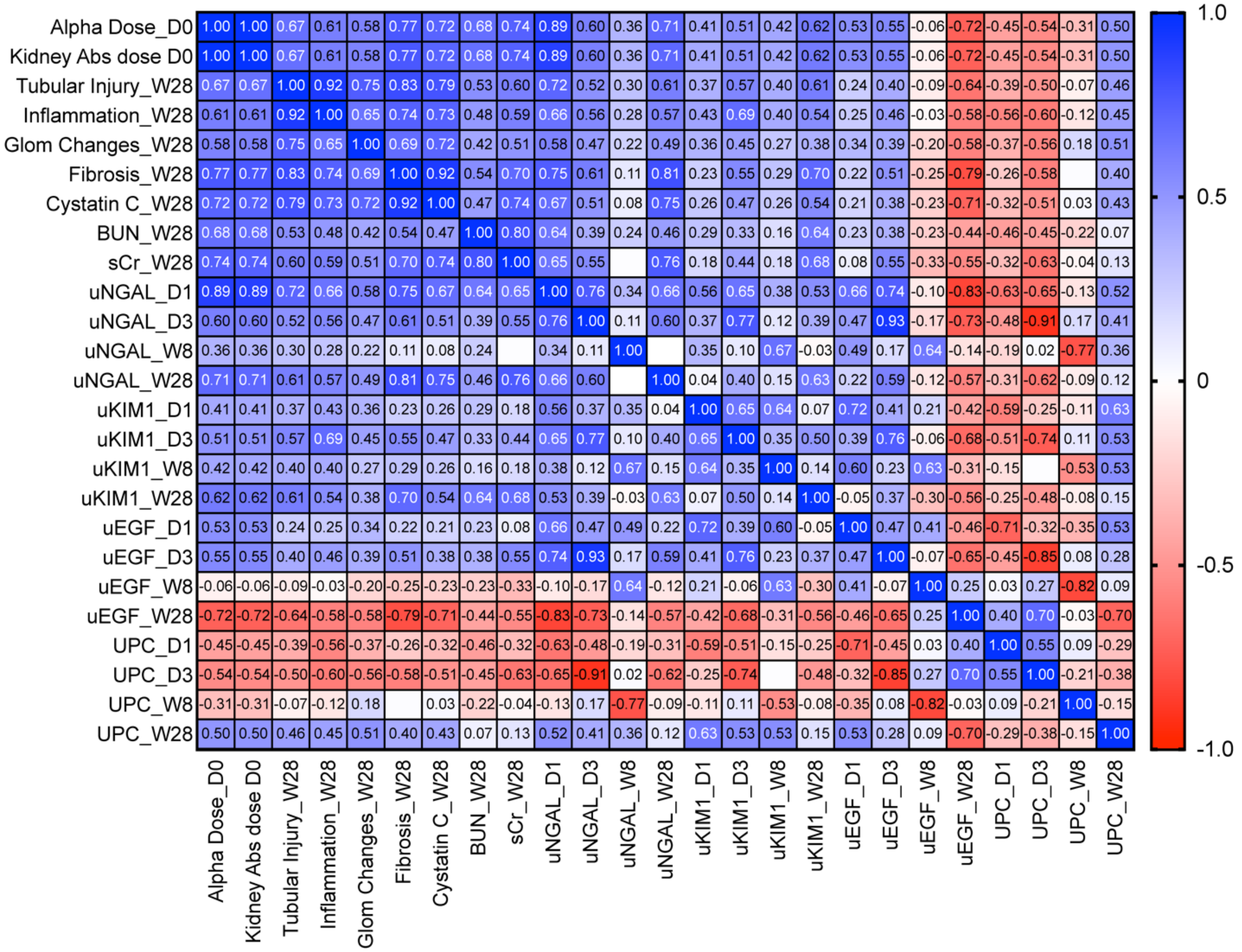
Alpha Dose_D0
Kidney Abs dose D0
Tubular Injury_W28
Inflammation_W28
Glom Changes_W28
Fibrosis_W28
Cystatin C_W28
BUN_W28
sCr_W28
uNGAL_D1
uNGAL_D3
uNGAL_W8
uNGAL_W28
uKIM1_D1
uKIM1_D3
uKIM1_W8
uKIM1_W28
uEGF_D1
uEGF_D3
uEGF_W8
uEGF_W28
UPC_D1
UPC_D3
UPC_W8
UPC_W28
1.0
0.5
0
-0.5
-1.0

**Fig Supp 3**

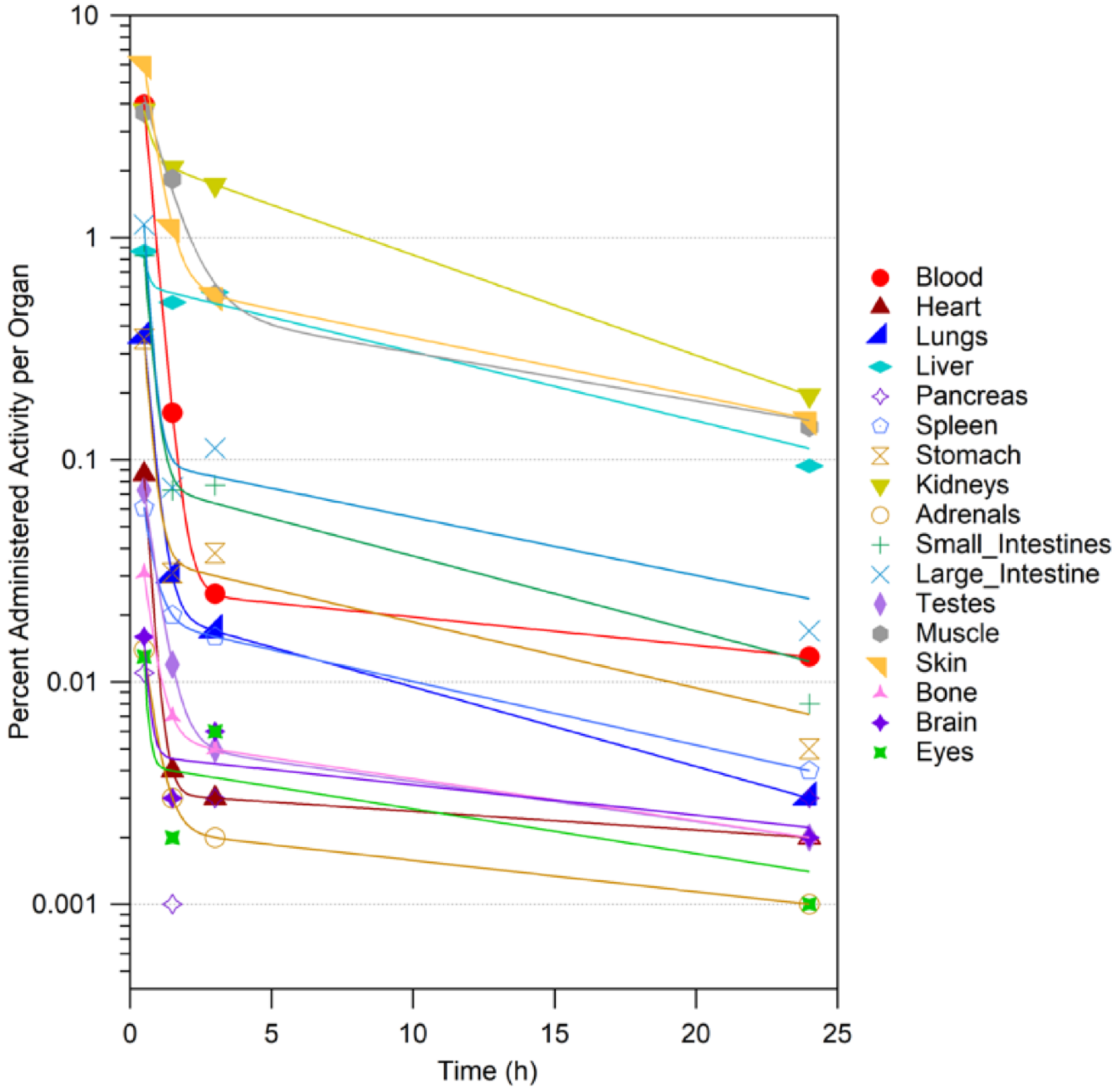

Percent Administered Activity per Organ
10
1
0.1
0.01
0.001
0
5
10
15
20
25
Time (h)
Blood
Heart
Lungs
Liver
Pancreas
Spleen
Stomach
Kidneys
Adrenals
Small_Intestines
Large_Intestine
Testes
Muscle
Skin
Bone
Brain
Eyes

**Fig Supp 4**

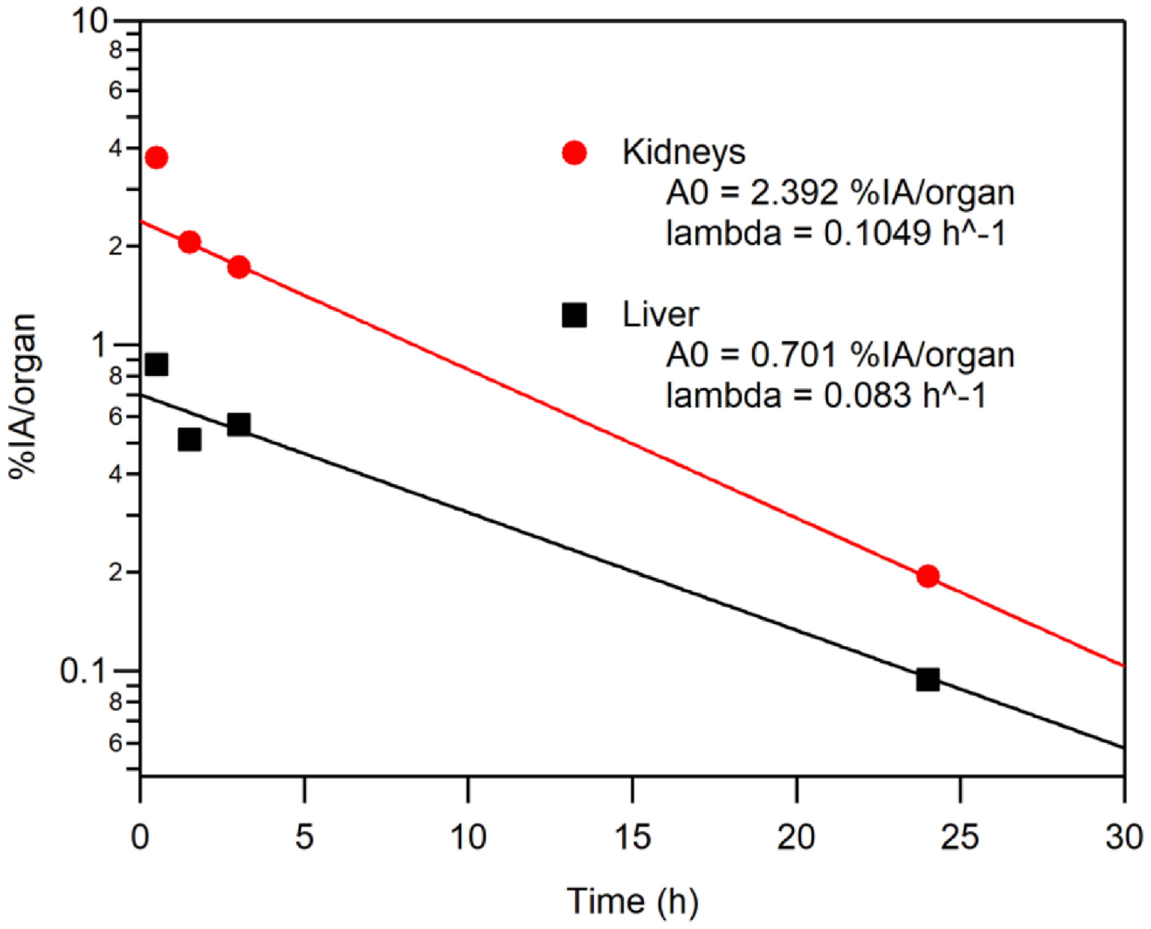

Kidneys
A0 = 2.392 %IA/organ
lambda = 0.1049 h^-1
Liver
A0 = 0.701 %IA/organ
lambda = 0.083 h^-1
%IA/organ
Time (h)
10
8
6
4
2
1
0.1
0
5
10
15
20
25
30
