## Supplemental Tables for "Pre-clinical Evaluation of Biomarkers for Early Detection of Nephrotoxicity Following Alpha-particle Radioligand Therapy"

**European Journal of Nuclear Medicine and Molecular Imaging**


Mengshi Li,[1*] Claudia Robles-Planells,[2*] Dijie Liu,[1] Stephen A. Graves,[3] Gabriela Vasquez-Martinez,[2] Gabriel Mayoral-Andrade,[2] Dongyoul Lee,[4] Prerna Rastogi,[5] Brenna M. Marks,[1] Edwin A. Sagastume,[1] Robert M. Weiss,[6] Sarah C. Linn-Peirano,[2,7] Frances L. Johnson,[1] Michael K. Schultz,[1,3,8#] Diana Zepeda-Orozco[2,9,10#]

[1]Viewpoint Molecular Targeting, Inc., Coralville, IA, USA

[2]Kidney and Urinary Tract Center, Abigail Wexner Research Institute at Nationwide Children's, Columbus, Ohio, USA

[3]Department of Radiology, The University of Iowa, Iowa City, IA, USA.

[4]Department of Physics and Chemistry, Korea Military Academy, Seoul, Republic of Korea

[5]Department of Pathology, The University of Iowa, Iowa City, IA, USA

[6]Department of Internal Medicine, The University of Iowa, Iowa City, IA, USA

[7]Department of Veterinary Biosciences, The Ohio State University College of Veterinary Medicine Columbus, OH

[8]Department of Pediatrics, The Ohio State University College of Medicine, Columbus, Ohio, USA.

[9]Division of Nephrology and Hypertension, Nationwide Children's Hospital, Columbus, Ohio, USA.

[10]Department of Radiation Oncology, Free Radical and Radiation Biology Program, The University of Iowa, Iowa City, IA, USA.

[*]Co-First Authors

[#]Co-Corresponsing Authors

Diana Zepeda-Orozco:

Schultz, Michael:

**Supplemental Table 1**. Scoring Criteria for Kidney Histological Changes

| Tubular Injury Scoring | | Glomerular Changes | |
|---|---|---|---|
| 0 | Absent | 0 | Absent |
| 1 | Mild (1-10%) | 1 | 1-10% |
| 2 | Moderate (11-25%) | 2 | 11-20% |
| 3 | Severe (26-50%) | 3 | 21-30% |
| 4 | Very Severe > 50% | 4 | 31% and greater |

| Tubulointerstitial Inflammation | | Interstitial Fibrosis | |
|---|---|---|---|
| 0 | Absent | 0 | Absent |
| 1 | Mild (1-10%) | 1 | Mild (1-10%) |
| 2 | Moderate (11-25%) | 2 | Moderate (11-25%) |
| 3 | Severe (26-50%) | 3 | Severe (26-50%) |
| 4 | Very Severe > 50% | 4 | Very Severe > 50% |

**Supplemental Table 2.** Ex vivo [$^{203}$Pb]Pb-MC1L biodistribution results for male CD-1 Elite mice (Units of percent injected dose per gram, corrected for [$^{203}$Pb] radioactive decay).

| $^{203}$Pb-VMT01 Percent injected dose per gram (%ID/g) | | | | | |
|---|---|---|---|---|---|
| **Tissue** | **0.5 h** | **1.5 h** | **3.0 h** | **24 h** | **Avg. Organ Mass (g)** |
| **Blood** | 1.37 | 0.06 | 0.01 | 0.02 | 3.000* |
| **Heart** | 0.68 | 0.03 | 0.03 | 0.06 | 0.132 |
| **Lungs** | 2.15 | 0.19 | 0.12 | 0.08 | 0.173 |
| **Liver** | 0.64 | 0.4 | 0.49 | 0.32 | 1.409 |
| **Pancreas** | 0.5 | 0.04 | 0.13 | 0.57 | 0.023 |
| **Spleen** | 0.63 | 0.22 | 0.19 | 0.21 | 0.101 |
| **Stomach** | 0.86 | 0.08 | 0.11 | 0.06 | 0.423 |
| **Kidneys** | 8.3 | 4.88 | 4.51 | 1.99 | 0.467 |
| **Adrenals** | 1.31 | 0.34 | 0.27 | 0.42 | 0.011 |
| **Small Intestines** | 0.64 | 0.06 | 0.07 | 0.03 | 1.334 |
| **Large Intestine** | 0.86 | 0.06 | 0.1 | 0.06 | 1.379 |
| **Testes** | 0.41 | 0.07 | 0.03 | 0.05 | 0.183 |
| **Muscle** | 0.34 | 0.18 | 0.06 | 0.06 | 11.250** |
| **Skin** | 1.12 | 0.22 | 0.12 | 0.13 | 5.625*** |
| **Bone** | 0.46 | 0.11 | 0.09 | 0.14 | 0.069 |
| **Brain** | 0.04 | 0.01 | 0.02 | 0.02 | 0.369 |
| **Eyes** | 0.4 | 0.08 | 0.23 | 0.12 | 0.032 |

* Assuming 8% of total body mass is blood.

** Assuming 30% of total body mass is muscle.

*** Assuming 15% of total body mass is skin.

**Supplemental Table 3.** Percent administered activity ([$^{212}$Pb]Pb-MC1L) per organ as a function of time, calculated from [$^{203}$Pb]Pb-MC1L ex vivo biodistribution data.

| $^{212}$Pb-VMT01 percent administered activity per (%AA/organ) | | | | |
|---|---|---|---|---|
| **Tissue** | **0.5 h** | **1.5 h** | **3.0 h** | **24 h** |
| **Blood** | 3.978 | 0.163 | 0.025 | 0.013 |
| **Heart** | 0.086 | 0.004 | 0.003 | 0.002 |
| **Lungs** | 0.359 | 0.030 | 0.017 | 0.003 |
| **Liver** | 0.867 | 0.511 | 0.568 | 0.094 |
| **Pancreas** | 0.011 | 0.001 | 0.003 | 0.003 |
| **Spleen** | 0.061 | 0.020 | 0.016 | 0.004 |
| **Stomach** | 0.350 | 0.031 | 0.038 | 0.005 |
| **Kidneys** | 3.750 | 2.066 | 1.731 | 0.195 |
| **Adrenals** | 0.014 | 0.003 | 0.002 | 0.001 |
| **Small Intestines** | 0.830 | 0.073 | 0.077 | 0.008 |
| **Large Intestine** | 1.147 | 0.075 | 0.113 | 0.017 |
| **Testes** | 0.073 | 0.012 | 0.005 | 0.002 |
| **Muscle** | 3.659 | 1.836 | 0.555 | 0.141 |
| **Skin** | 6.098 | 1.122 | 0.555 | 0.153 |
| **Bone** | 0.031 | 0.007 | 0.005 | 0.002 |
| **Brain** | 0.016 | 0.003 | 0.006 | 0.002 |
| **Eyes** | 0.013 | 0.002 | 0.006 | 0.001 |

**Supplemental Table 4.** Bi-exponential fit parameters, and time-integrated activity coefficients for the two integration methods. $\tau 1$: Time-integrated activity coefficient calculated from the bi-exponential fit. $\tau 2$: time integrated activity coefficient calculated from trapezoidal integration assuming constant activity from 0 to 0.5 h, and physical decay beyond the terminal timepoint.

| Fit coefficients: | $A_1$ | $\lambda_1$ | $A_2$ | $\lambda_2$ | $\tau_1$ | $\tau_2$ | $\tau_{avg}$ |
|---|---|---|---|---|---|---|---|
| Blood | 21.164 | 3.356 | 0.026 | 0.029 | 0.0720 | 0.0478 | 0.0599 |
| Heart | 0.789 | 4.508 | 0.003 | 0.019 | 0.0034 | 0.0017 | 0.0026 |
| Lungs | 1.890 | 3.441 | 0.022 | 0.082 | 0.0081 | 0.0066 | 0.0074 |
| Liver | 12.640 | 7.763 | 0.063 | 0.072 | 0.0250 | 0.1033 | 0.0642 |
| Pancreas | * | * | * | * | * | 0.0011 | 0.0011 |
| Spleen | 0.178 | 2.878 | 0.019 | 0.066 | 0.0036 | 0.0038 | 0.0037 |
| Stomach | 2.598 | 4.226 | 0.037 | 0.068 | 0.0115 | 0.0096 | 0.0106 |
| Kidneys | 8.897 | 3.553 | 2.364 | 0.104 | 0.2524 | 0.3084 | 0.2804 |
| Adrenals | 0.043 | 2.555 | 0.002 | 0.033 | 0.0008 | 0.0007 | 0.0008 |
| Small Intestines | 5.745 | 4.067 | 0.080 | 0.078 | 0.0245 | 0.0200 | 0.0222 |
| Large Intestine | 11.872 | 4.853 | 0.101 | 0.060 | 0.0412 | 0.0297 | 0.0354 |
| Testes | 0.212 | 2.281 | 0.005 | 0.041 | 0.0022 | 0.0019 | 0.0021 |
| Muscle | 7.112 | 1.191 | 0.511 | 0.051 | 0.1600 | 0.1585 | 0.1593 |
| Skin | 17.542 | 2.330 | 0.645 | 0.060 | 0.1829 | 0.1771 | 0.1800 |
| Bone | 0.099 | 2.709 | 0.006 | 0.043 | 0.0017 | 0.0015 | 0.0016 |
| Brain | 0.243 | 6.130 | 0.005 | 0.031 | 0.0019 | 0.0013 | 0.0016 |
| Eyes | 0.462 | 7.928 | 0.004 | 0.046 | 0.0015 | 0.0011 | 0.0013 |

*Regression not well defined for Pancreas due to increase in estimated tissue concentration after 1.5 hours.

**Supplemental Table 5.** Summary of mouse and human organ masses and organ [$^{212}$Pb]Pb-MC1L time-integrated activity coefficients.

| Tissue | $m_{mouse}$ (g) | $\tau_{mouse}$ (MBq*h/MBq) | $m_{human}$ (g) | $\tau_{human}$ (MBq*h/MBq) |
|---|---|---|---|---|
| **Blood** | 3.000 | 5.99E-02 | 6364 | 2.42E-01 |
| **Heart** | 0.132 | 2.55E-03 | 330 | 8.61E-03 |
| **Lungs** | 0.173 | 7.38E-03 | 1200 | 8.14E-02 |
| **Liver** | 1.409 | 6.42E-02 | 1800 | 1.01E-01 |
| **Pancreas** | 0.023 | 1.13E-03 | 140 | 3.46E-03 |
| **Spleen** | 0.101 | 3.67E-03 | 150 | 4.30E-03 |
| **Stomach** | 0.423 | 1.06E-02 | 250 | 8.56E-03 |
| **Kidneys** | 0.467 | 2.80E-01 | 310 | 3.22E-01 |
| **Adrenals** | 0.011 | 7.77E-04 | 14 | 8.00E-04 |
| **Small Intestines** | 1.334 | 2.22E-02 | 350 | 8.41E-03 |
| **Large Intestine** | 1.379 | 3.54E-02 | 225 | 8.54E-03 |
| **Testes** | 0.183 | 2.06E-03 | 35 | 4.51E-04 |
| **Muscle** | 11.250 | 1.59E-01 | 29200 | 4.24E-01 |
| **Skin** | 5.625 | 1.80E-01 | 3600 | 1.32E-01 |
| **Bone** | 0.069 | 1.58E-03 | 4400 | 8.61E-02 |
| **Brain** | 0.369 | 1.59E-03 | 1450 | 5.36E-03 |
| **Eyes** | 0.032 | 1.28E-03 | 15 | 6.86E-04 |

**Supplemental Table 6.** Murine absorbed dose estimates by tissue for [$^{212}$Pb]Pb-MC1L, generated using OLINDA 2.2.

| Tissue | Dose per administered activity (mGy/MBq) Alpha | Beta | Gamma | Total |
|---|---|---|---|---|
| **Brain** | 2.05E+01 | 7.46E+00 | 4.14E-01 | 2.84E+01 |
| **Large Intestine** | 1.67E+02 | 2.14E+01 | 7.75E-01 | 1.89E+02 |
| **Small Intestine** | 1.26E+02 | 1.76E+01 | 1.40E+00 | 1.45E+02 |
| **Stomach** | 1.64E+02 | 1.94E+01 | 3.48E-01 | 1.84E+02 |
| **Heart** | 1.37E+02 | 2.03E+01 | 1.15E+00 | 1.59E+02 |
| **Kidneys** | 2.76E+03 | 2.34E+02 | 3.18E+00 | 2.99E+03 |
| **Liver** | 2.09E+02 | 2.87E+01 | 1.41E+00 | 2.39E+02 |
| **Lungs** | 1.99E+02 | 1.52E+01 | 5.18E-01 | 2.14E+02 |
| **Pancreas** | 2.16E+02 | 2.89E+02 | 9.81E+00 | 5.15E+02 |
| **Skeleton** | 1.29E+02 | 5.30E+01 | 6.23E+00 | 1.88E+02 |
| **Spleen** | 1.06E+02 | 2.10E+01 | 1.23E+00 | 1.29E+02 |
| **Testes** | 4.98E+01 | 9.81E+00 | 5.21E-01 | 6.01E+01 |
| **Thyroid** | 4.91E+01 | 5.68E+00 | 4.97E-01 | 5.53E+01 |
| **Total Body** | 4.91E+01 | 1.14E+01 | 4.87E-01 | 1.14E+02 |
| **Blood*** | 8.98E+01 | - | - | 8.98E+01 |
| **Adrenals*** | 3.13E+02 | - | - | 3.13E+02 |
| **Muscle*** | 6.37E+01 | - | - | 6.37E+01 |
| **Skin*** | 1.44E+02 | - | - | 1.44E+02 |
| **Eyes*** | 1.78E+02 | - | - | 1.78E+02 |

* Blood, adrenals, muscle, skin, and eyes are not implemented as source/target tissues in the OLINDA 2.2 murine models, and therefore dose to these tissues was calculated assuming local alpha particle energy deposition, with an average alpha energy of 7.8063 MeV per [$^{212}$Pb] decay. Beta and gamma self-dose contributions were not calculated for these tissues.

**Supplemental Table 7.** Estimated human dosimetry for [212Pb]Pb-MC1L.

| Tissue | Alpha (mGy/MBq) | Beta (mGy/MBq) | Gamma (mGy/MBq) | Total (mGy/MBq) |
|---|---|---|---|---|
| **Adrenals** | 2.62E-01 | 4.54E-02 | 3.26E-02 | 3.40E-01 |
| **Brain** | 1.69E-02 | 1.89E-03 | 2.07E-03 | 2.09E-02 |
| **Esophagus** | 4.70E-02 | 5.32E-03 | 7.24E-03 | 5.96E-02 |
| **Eyes** | 2.10E-01 | 2.17E-02 | 3.37E-03 | 2.35E-01 |
| **Gallbladder** | 4.70E-02 | 6.04E-03 | 1.14E-02 | 6.44E-02 |
| **Left Colon** | 4.70E-02 | 1.52E-02 | 9.92E-03 | 7.21E-02 |
| **Small Intestine** | 4.70E-02 | 1.16E-02 | 7.98E-03 | 6.65E-02 |
| **Stomach** | 4.70E-02 | 1.44E-02 | 8.87E-03 | 7.03E-02 |
| **Right Colon** | 4.70E-02 | 1.52E-02 | 8.96E-03 | 7.11E-02 |
| **Rectum** | 4.70E-02 | 5.29E-03 | 5.60E-03 | 5.78E-02 |
| **Heart** | 1.20E-01 | 1.36E-02 | 7.77E-03 | 1.41E-01 |
| **Kidneys** | **4.76E+00** | **5.17E-01** | **5.91E-02** | **5.34E+00** |
| **Liver** | 2.58E-01 | 2.90E-02 | 1.37E-02 | 3.01E-01 |
| **Lungs** | 3.11E-01 | 3.33E-02 | 7.02E-03 | 3.51E-01 |
| **Pancreas** | 1.13E-01 | 1.23E-02 | 1.03E-02 | 1.36E-01 |
| **Prostate** | 4.70E-02 | 5.29E-03 | 5.69E-03 | 5.79E-02 |
| **Salivary** | 4.70E-02 | 5.29E-03 | 3.60E-03 | 5.59E-02 |
| **Red Marrow** | **2.43E-01** | **1.32E-02** | **6.39E-03** | **2.63E-01** |
| **Osteogenic Cells** | 1.17E+00 | 2.60E-02 | 6.11E-03 | 1.20E+00 |
| **Spleen** | 1.31E-01 | 1.54E-02 | 1.38E-02 | 1.61E-01 |
| **Testes** | 5.91E-02 | 6.23E-03 | 3.50E-03 | 6.87E-02 |
| **Thymus** | 4.70E-02 | 5.41E-03 | 5.68E-03 | 5.80E-02 |
| **Thyroid** | 4.70E-02 | 5.29E-03 | 4.78E-03 | 5.71E-02 |
| **Urinary Bladder Wall** | 4.70E-02 | 5.29E-03 | 4.91E-03 | 5.72E-02 |
| **Total Body** | 8.84E-02 | 1.02E-02 | 4.66E-03 | 1.03E-01 |
